# A genome-wide CRISPR-Cas9 screen identifies TGN46 as a host determinant for H-1PV susceptibility in pancreatic adenocarcinoma

**DOI:** 10.64898/2026.03.06.710102

**Authors:** Guillaume Labrousse, Nina Suarez, Adèle Nevot, Naïma Hanoun, Nelson Dusetti, Louis Buscail, Claudio Murgia, Pierre Cordelier

## Abstract

Pancreatic ductal adenocarcinoma (PDAC) remains a highly lethal malignancy with limited response to current therapeutic strategies. Oncolytic viruses such as the rat protoparvovirus H-1PV represent promising alternatives; however, their clinical development is hindered by an incomplete understanding of virus-host interactions that determine tumor susceptibility. Here, we conducted a genome-scale CRISPR-Cas9 knockout screen in primary PDAC cultures to identify host determinants of viral infection and cytotoxicity. The screen confirmed the central role of the sialylation pathway in mediating susceptibility to H-1PV and uncovered several additional genes involved in vesicular trafficking and protein recycling. Among these candidates, *TGOLN2*, encoding the transmembrane protein TGN46, emerged as a critical host factor. Functional analyses demonstrate that TGN46 supports efficient H-1PV infection, replication, and virus-induced cytotoxicity in pancreatic cancer cells. TGN46 colocalizes with viral particles at the cell surface and facilitates their internalization through dynamin-dependent endocytosis. Molecular studies further indicate that the luminal N-terminal domain of TGN46 interacts with the viral capsid. *In vivo*, *TGOLN2* expression in essential to H-1PV-mediated antitumor activity in experimental models. Collectively, these findings identify TGN46 as a membrane-associated entry factor required for optimal H-1PV infection in PDAC cells. This work refines the mechanistic understanding of H-1PV tropism and provides a rationale for exploring host determinants of viral susceptibility as candidate biomarkers to guide parvovirus-based virotherapy.

## Introduction

Pancreatic adenocarcinoma (PDAC) is notoriously resistant to therapy and difficult to cure, with a 5-year survival rate of patients less than 11%^1,2^. This poor prognosis is primarily due to the limited efficacy of current treatments, despite recent advances^3^. Most existing options, primarily chemotherapies, have shown modest benefits. In contrast, oncolytic viruses (OVs) are emerging as a promising therapeutic alternative. OVs are naturally occurring or engineered viruses that selectively infect and kill cancer cells while sparing normal tissues. They replicate within tumor cells, causing direct oncolysis and releasing tumor antigens, which can stimulate systemic anti-tumor immunity^4^. Notable example is Talimogene laherparepvec (T-VEC), an engineered herpes simplex virus approved for melanoma treatment, which has demonstrated clinical efficacy^5^. Various OVs have been investigated in preclinical and clinical settings for their ability to target and eliminate PDAC cells^6^. However, the varying efficacy of OVs in PDAC including the limited success of T-VEC^7^, highlights the critical need to understand the molecular and cellular mechanisms underlying oncolytic activity. In particular, identifying host factors in tumor cells may help refine patient selection and optimize therapeutic outcomes with virotherapy.

Among the OV strains currently under investigation for PDAC, the rat H-1 protoparvovirus (H-1PV) displays particularly promising properties. H-1PV is not a human pathogen and is not associated with any known human disease. It demonstrates natural tropism and strong oncolytic activity across various cancer cell types, both *in vitro* and *in vivo* ^8^. Moreover, it is well-tolerated in patients, with no severe adverse effects directly attributed to the virus, in phase I/IIa and phase II clinical trials in glioblastoma ^9^ and PDAC ^10,11^. These findings support the potential of H-1PV as a viable candidate for oncolytic virotherapy in PDAC. H-1PV is a small, 25 nm encapsidated virus that carries a 5 kb linear single-stranded DNA genome encoding four proteins from two genes. The early P4 promoter drives expression of the non-structural (NS) proteins NS1 and NS2. NS1 is essential for viral genome replication in the host cell nucleus and activates the late P38 promoter, which controls the expression of the viral capsid proteins VP1 and VP2. These capsid proteins assemble into new virions following viral DNA packaging^12^. The oncolytic activity of H-1PV primarily relies on NS1-induced production of reactive oxygen species (ROS) and subsequent cell death resulting from DNA damage^13^.

Significant efforts have been made to better understand H-1PV and its interaction with host cells. An emerging model of H-1PV entry highlights the role of the laminin γ1 chain in anchoring viral particles to the extracellular matrix (ECM)^14^. Galectin-1 (Gal-1) is also implicated in this process, cooperating with laminin during viral entry, although it does not contribute to the initial attachment of viral particles to the cell surface^14,15^. Sialic acid (SIA) metabolism probably plays a central role in this mechanism, as H-1PV binding to laminin γ1 depends on specific SIA glycosylation motifs. Galectins can further interact with sialylated proteins to modulate viral infection, depending on the particular sialylation pattern^16^. Following attachment, H-1PV enters cells via clathrin-mediated endocytosis (CME)^17^, eventually escaping from the endosomal compartment^18^. However, despite being discovered more than 60 years ago^19^, a specific cellular receptor for H-1PV has yet to be identified.

In this study, we aimed to identify novel host genes involved in H-1PV infection of PDAC cells, with a particular focus on discovering candidates that may serve as potential viral receptors. We performed a genome-scale CRISPR/Cas9 knockout (KO) screen in primary PDAC cultures with intermediate sensitivity to viral infection and confirmed the essential role of the protein sialylation pathway in H-1PV susceptibility. In parallel, we identified several host genes associated with vesicular recycling, retrograde transport, and protein secretion. Among these, we focused on *TGOLN2* (Trans-Golgi Network Protein 2), which encodes TGN46, a glycosylated transmembrane protein that cycles rapidly between the cell surface and the trans-Golgi network (TGN)^20^. We demonstrated that H-1PV critically depends on TGN46 for efficient infection, replication, and cytotoxicity in primary PDAC cells. Further molecular analyses indicate that TGN46 colocalizes with H-1PV at the cell surface and facilitates rapid viral internalization in a dynamin-dependent manner. We also provide evidence that the N-terminal luminal domain of TGN46 interacts with the H-1PV capsid, in a process influenced by sialylation. Moreover, TGN46 expression associates with H-1PV permissiveness and antitumor activity in pancreatic cancer models, both *in vitro* and *in vivo*. Together, these findings identify TGN46 as a membrane-associated entry factor required for efficient H-1PV infection and refine the current understanding of parvovirus tropism. This work establishes a mechanistic basis for further exploration of host determinants as potential biomarkers of response to parvovirus-based virotherapy.

## Materials and Methods

### Cell culture and treatment

HeLa S3 (ATCC CCL-2™), HEK-293T (ATCC CRL-3216™), and human newborn kidney cells NB-324K (kindly provided by Dr. Jean Rommelaere, DKFZ, Heidelberg, Germany) were cultured in Dulbecco’s Modified Eagle Medium (DMEM, 4.5 g/L glucose; Sigma-Aldrich, D6429) supplemented with 10% fetal calf serum (FCS; Gibco, 10270-106) and 1× antibiotic-antimycotic solution (Gibco, 15240-062). Human primary pancreatic cancer cells were maintained as previously described^21^. All cells were cultured at 37°C in a humidified atmosphere containing 5% CO₂. Doxycycline (Dox; Sigma-Aldrich, D9891) was dissolved in PBS at a stock concentration of 0.5 mM and used at a final concentration of 0.5 µM. For inducible overexpression experiments, cells were pre-treated with doxycycline at least 24 hours before seeding and continuously exposed throughout the experiment. Dynasore (Tocris Bioscience, 2897), a dynamin inhibitor, was dissolved in DMSO at a stock concentration of 25 mM and used at a final concentration of 100 µM. Cells were pre-treated for 2 hours prior to experimentation and maintained under treatment during the assay. Transferrin from human serum conjugated to Alexa Fluor™ 647 (TFn-647; Fisher Scientific, 11550766) was reconstituted in deionized water at a stock concentration of 5 mg/mL and applied to cells at a final concentration of 25 µg/mL for 5 minutes prior to fixation. For antibiotic selection, blasticidin (InvivoGen, ant-bl) was used at 0.5 µg/mL, puromycin (InvivoGen, ant-pr) at 1 µg/mL, and ouabain (Sigma-Aldrich, O3125) at 0.5 µM. Selection was carried out for a minimum of one week.

### Plasmid amplification and transfection

DNA plasmid constructs used in this study were transformed and amplified in E. coli Stbl3™ competent cells (Invitrogen - C7373-03). Bacteria were plated on LB agar supplemented with the appropriate antibiotic, and individual colonies were expanded for plasmid extraction. Initial plasmid preparations were performed using the ZR Plasmid Miniprep Classic kit (Zymo Research, D4016). Clones were verified by restriction enzyme digestion and Sanger sequencing (Eurofins Genomics, LightRun). For large-scale preparations, endotoxin-free plasmid DNA was obtained using the NucleoBond Xtra Maxi kit (Macherey-Nagel, 740414.50). At each step, plasmid DNA concentration and purity were assessed using a NanoDrop spectrophotometer (ND-1000, Thermo Scientific). Cell transfections were performed using JetPRIME transfection reagent (Polyplus, 101000046) according to the manufacturer’s instructions, using a DNA:JetPRIME ratio of 1:2 (µg:µL).

### Virus Production

The initial stock of H-1 protoparvovirus (H-1PV) was kindly provided by Dr. Jean Rommelaere (DKFZ, Heidelberg, Germany). For virus amplification, 1.5 × 10⁶ NB-324K cells were seeded in T175 flasks and cultured until reaching approximately 30% confluency. Cells were then infected with H-1PV at a multiplicity of infection (MOI) of 10⁻² in 3 mL of infection medium (IM) consisting of Opti-MEM (Gibco, 31985-047) supplemented with 1% HEPES (Gibco, 15630-056) for 1 hour at 37 °C. After the adsorption period, the medium was replaced with 20 mL of fresh Opti-MEM, and cells were incubated at 37 °C for 4 days until the appearance of cytopathic effects (CPE). Cells were harvested by scraping into a minimal volume of fresh Opti-MEM. Virions were released by three freeze–thaw cycles using liquid nitrogen, followed by sonication on ice at 25 W for 30 seconds. Cellular debris were removed by centrifugation at 1,800 rpm for 15 minutes, and the supernatant was filtered through a 0.2 μm membrane (ClearLine, 146560). Viral stocks were aliquoted and stored at −80 °C. Viral titers were determined by plaque assay on NB-324K cells as previously described^22^.

### Lentivirus production and cell transduction

On day 1, for each lentiviral production, 4.5 × 10⁶ HEK-293T cells were seeded in a 100 mm dish with 10 mL of complete DMEM. On day 2, HEK-293T cells were co-transfected with 6 µg of lentiviral plasmid, 4 µg of the psPAX2 packaging plasmid (Addgene #12260), and 1.5 µg of the pMD2.G envelope plasmid (Addgene #12259) using JetPRIME reagent. Four hours post-transfection, the medium was replaced with 10 mL of Opti-MEM. On day 3, target cells were seeded in 6-well plates to reach 30–40% confluency by the next day. On day 4, HEK-293T supernatants were collected in 50 mL tubes, centrifuged for 5 minutes at 1800 rpm to remove cell debris, and the supernatant was filtered through a 0.45 µm filter (Millex-HP, SLHP033RS). Lentiviral supernatants were concentrated using Vivaspin 20 columns (100 kDa MWCO PES; Sartorius, VS2042) by centrifugation at 2500 rpm for 20–30 minutes. Target cells were transduced with the lentiviral vectors in 2 mL of Opti-MEM supplemented with protamine sulfate at a final concentration of 10 µg/mL. Forty-eight hours post-transduction, cells were expanded, and appropriate antibiotic selection was initiated the following day. Selection was maintained for at least one week, and cells were passaged at least twice before experimental use.

### CRISPR knock-out screen

The CRISPR knock-out (KO) screen methodology was adapted from^23^. Briefly, PDAC087T cells were first transduced with lentiCas9-Blast (Addgene #52962), enabling constitutive expression of the Cas9 enzyme and resistance to blasticidin. After antibiotic selection and expansion, PDAC087T-Cas9 cells were seeded in five T175 flasks at 1 × 10⁷ cells per flask and transduced with the GeckoV2 pooled gRNA libraries A and B, inserted in the lentiGuide-Puro plasmid (Addgene #1000000049), at a multiplicity of infection (MOI) of ∼0.3. This low MOI maximizes the probability of integrating a single sgRNA per cell while maintaining library complexity. Following one week of puromycin selection, 7.5 × 10⁶ PDAC087T-Cas9-GeckoV2 cells were seeded in seven T175 flasks per condition (>5 × 10⁷ cells total). The following day, cells were incubated for 1 h at 37 °C with 5 mL infection medium (IM), with or without H-1PV at MOI 10 (a dose previously shown to induce >90% cell death in PDAC087T cells within 48 h). After infection, cells were washed with PBS and incubated in 20 mL of complete DMEM at 37 °C. Non-infected cells were passaged at a 1:10 dilution upon reaching confluency. After one week, dead and floating cells were removed by repeated PBS washes. Adherent cells from each condition were detached by trypsinization, centrifuged, resuspended in complete medium, and pooled into one T175 flask per condition. Two days later, cells were harvested again, resuspended in PBS, counted, and genomic DNA was extracted using the DNeasy Blood & Tissue Kit (Qiagen #69504; one column per 5 × 10⁶ cells, with elution in minimal volume). For each condition, sgRNA loci were amplified using NGS-grade barcoded PCR primers (Eurogentec, Table 1). Amplicons were size-separated by electrophoresis on 2% agarose gel, purified with the NucleoSpin Gel and PCR Clean-up Kit (Macherey-Nagel #740609.250), and further cleaned using SPRIselect Beads (Beckman Coulter #B24965AA). DNA fragment size was monitored at each step using a Fragment Analyzer (Agilent). Once quality was confirmed, barcoded amplicons were sequenced on the NextSeq 550 platform (Illumina). Sequencing data were demultiplexed using the bcl2fastq algorithm (Illumina) and analyzed for sgRNA enrichment using MAGeCK (Model-based Analysis of Genome-wide CRISPR-Cas9 Knockout)^24^.

### Generation of knock-out cell lines

To generate knock-out (KO) cell lines for individual genes of interest, sgRNA protospacer sequences targeting specific loci were ordered as complementary oligonucleotides. Each oligo pair was phosphorylated using T4 Polynucleotide Kinase (NEB #M0201) and annealed in a T100 Thermal Cycler (Bio-Rad #1861096) by heating at 95 °C for 5 min, followed by a gradual temperature ramp down at 0.1 °C/sec to 25 °C. Subcloning of the annealed protospacer sequences was performed by Golden Gate assembly. Appropriate cohesive ends were designed to enable ligation of the protospacer DNA into either the lentiGuide-Puro plasmid (Addgene #52963) or the eSpCas9 (1.1) No FLAG ATP1 A1G2 Dual sgRNA plasmid (Addgene #86612), previously digested with BsmBI (NEB #R0739) or BbsI (NEB #R3539), respectively. For validation of sgRNAs identified in the CRISPR KO screen, PDAC087T-Cas9 cells were thawed and seeded in 6-well plates at a density of 1.5 × 10⁵ cells per well. The following day, each well was transduced with a different lentiGuide-Puro construct carrying one sgRNA to validate. Cells were then selected with puromycin prior to downstream use. The sequences of the sgRNAs used are listed in Table 2. To generate *TGOLN2*-KO cells while avoiding constitutive expression of Cas9 and sgRNA, PDAC087T and HeLa S3 cells were seeded in 6-well plates at 1.5 × 10⁵ cells per well and transfected the following day with 3 µg of the eSpCas9 (1.1) No FLAG ATP1A1 G2 Dual sgRNA plasmid. This construct expresses Cas9 along with a sgRNA targeting exon 3 of *TGOLN2*. To enrich for successfully edited cells, the plasmid also contains a second sgRNA targeting exon 4 of the ATP1A1 gene, an essential gene whose disruption confers resistance to ouabain, a cytotoxic drug. This strategy selectively enriches cells that have undergone mutagenic repair at the ATP1A1 locus without impairing its function^25^. For generation of TGN46-WT control cells undergoing the same transfection and selection process, a similar construct expressing only the ATP1A1-targeting sgRNA was transfected in parallel. Five days post-transfection, ouabain was added to the culture medium. After one week of recovery, *TGOLN2* KO efficiency was assessed by Western blot and immunofluorescence.

### TGOLN2 cDNA cloning

Total mRNA from PDAC087T cells was extracted using the RNeasy Plus Mini Kit (Qiagen #74134) and reverse-transcribed into complementary DNA (cDNA) with the RevertAid Kit (Thermo Fisher #K1691). The resulting cDNA was used as a template for high-fidelity PCR amplification of the open reading frame (ORF) corresponding to the main *TGOLN2*-201 spliced variant (Ensembl transcript ID: ENST00000377386.8), which encodes the 437-amino acid TGN46 protein (6050 bp). PCR amplification was performed using PrimeSTAR Max DNA Polymerase (Takara #R045A). To facilitate downstream cloning, gene-specific primers containing restriction sites were used: an AvrII site at the 5’ end and a MluI site at the 3’ end. The sequences were as follows: Forward: 5′-TTTACGCGTttaggacttctggtccaaac-3′, Reverse: 5′-TTTCCTAGGgccaccatgcggttcgtggttgcctt-3. The PCR product was purified using the NucleoSpin Gel and PCR Clean-up Kit (Macherey-Nagel), digested for 1 h at 37 °C with AvrII and MluI restriction enzymes (NEB #R3198, #R0174), and re-purified before ligation into a similarly digested third-generation lentiviral vector, pLIX402 (Addgene #41394), to generate the pTRE-*TGOLN2* construct. This all-in-one plasmid contains the complete Tet-On system, including a constitutive expression cassette encoding the rtTA-3 transactivator, followed by a 2A self-cleaving peptide and the puromycin resistance gene. Upon doxycycline (dox) treatment, rtTA-3 binds to the TRE (tetracycline-responsive element) promoter, thereby inducing the expression of TGN46.

### TGOLN2 mutagenesis and fusion with EGFP and FLAG tags

TGN46 deletion constructs, TGN46-ΔI, TGN46-ΔII, and TGN46-ΔIII, were generated by high-fidelity PCR using the pTRE-*TGOLN2* plasmid as a template. Each deletion targeted specific domains of the protein as described in Illustration 1, and the oligonucleotides used for amplification are listed in Table 3. To facilitate detection and localization studies, EGFP or FLAG (DYKDDDDK) tags were fused either to the N- or C-terminus of TGN46. Importantly, N-terminal tags were inserted immediately downstream of the signal peptide sequence to preserve the protein’s tethering to the secretory pathway. The tagged and truncated TGN46 variants were subcloned into the pLIX402 lentiviral vector (Addgene #41394) using standard molecular cloning techniques detailed in Illustration 1. Lentiviral particles were produced using these constructs, and PDAC087T-*TGOLN2*-KO cells were transduced accordingly. After puromycin selection, cells expressing the tagged or truncated TGN46 variants were used in rescue experiments.

### Protein extraction and Western Blot analysis

For protein extraction, cultured cells grown in 100 mm dishes were washed with ice-cold PBS after removal of the culture medium. PBS was then aspirated, and cells were lysed directly on ice for 10 minutes with 200 µL of RIPA Buffer II, pH 8.0 (Bio Basic – RB4476) supplemented with protease inhibitor cocktail (Sigma-Aldrich – P8340). Lysates were scraped on ice and collected in 1.5 mL microcentrifuge tubes. Cell lysates were sonicated (e.g., 3 cycles of 5 seconds ON / 10 seconds OFF at 30% amplitude using a probe sonicator; adjust based on equipment), followed by centrifugation at 13,000 × g for 10 minutes at 4 °C. The clarified supernatants were transferred to fresh tubes, and protein concentrations were determined using the Bradford reagent (SERVA – 39222.03). A fixed amount of protein (30 to 80 µg per sample) was denatured in Laemmli buffer by heating at 95 °C for 5 minutes, then immediately loaded on 10–12% SDS-polyacrylamide gels (Sigma-Aldrich – A3699). Electrophoresis was performed using TG-SDS running buffer (Euromedex – EU0510). Proteins were transferred to nitrocellulose membranes using the Trans-Blot® Turbo™ RTA Transfer Kit and Transfer System (Bio-Rad – 1704271 and 1704150EDU). Transfer efficiency was verified by Ponceau S staining. Membranes were blocked for 30 minutes at room temperature (RT) on a rocking platform in TBS-T (TBS with 0.1% Tween-20; Sigma-Aldrich – P1379) containing 5% non-fat dry milk (Millipore – 70166). After blocking, membranes were incubated overnight at 4 °C with primary antibodies diluted in TBS-T + 1% milk, according to the manufacturers’ recommendations. The next day, membranes were washed three times for 10 minutes at RT in TBS-T, then incubated for 1 hour at RT with secondary HRP-conjugated antibodies diluted in TBS-T + 1% milk. After washing again (3×10 min, RT), detection was performed using the Clarity Western ECL substrate (Bio-Rad – 170-5061), and signal acquisition was carried out with the ChemiDoc™ XRS+ System and Image Lab™ Software (Bio-Rad – 1708265). All antibodies used, along with their sources and working dilutions, are listed in Table 4. Rabbit antisera against H-1PV NS1 and VP were generously provided by Dr. Jürg Nüesch (DKFZ, Heidelberg, Germany)^26^.

### Viability assays

Cells were seeded in white-bottom 96-well plates (Greiner Bio-One – 675075) at a density of 1 × 10⁴ cells per well in 100 µL of complete medium, in the presence or absence of dox. The following day, cells were incubated for 1 hour at 37 °C with 25 µL of infection medium containing or not H-1PV, applied across 10 different MOIs using a 1:2 serial dilution series starting from MOI 100 down to MOI 0.19. Following incubation, wells were washed once with PBS, and 100 µL of fresh complete medium (± dox) was added. At 48 hours post-infection, 100 µL of CellTiter-Glo® Luminescent Cell Viability reagent (Promega – G7572) was added per well. Plates were mixed for 2 minutes on an orbital shaker to ensure complete cell lysis and reagent mixing. Luminescence was measured as Relative Luminescence Units (RLU) using a TriStar² LB 942 plate reader (Berthold Technologies). For normalization, non-infected control wells were used to define 100% viability, and values from infected conditions were expressed as percentages relative to this reference.

### Cell counting

Cells were seeded in 24-well plates at a density of 4.5 × 10⁴ cells per well in 0.5 mL of complete DMEM, with or without dox. For each condition, four replicate wells were prepared, and a separate plate was used for each time point (4 h, 24 h, 48 h, and 72 h). The following day, cells were incubated for 1 hour at 37 °C with 250 µL of infection medium (IM), in the presence or absence of H-1PV, at MOIs of 2.5 or 10. After incubation, wells were washed three times with PBS, and 0.5 mL of fresh complete medium (± dox) was added. For cell counting, 200 µL of the cell suspension was diluted in 20 mL of ISOTON II diluent (Beckman Coulter – 8448011) and analyzed using a Coulter Z1 Particle Counter (Beckman Coulter).

### Viral genome quantification by RT(q)PCR

To assess virus production by infected cells, cells were seeded in 24-well plates at a density of 4.5 × 10⁴ cells per well in 0.5 mL of complete DMEM, with or without doxycycline (dox). For each condition, four replicate wells were prepared, and a separate plate was used for each time point (4 h, 24 h, 48 h, and 72 h). The following day, cells were incubated for 1 hour at 37°C with 250 µL of infection medium, in the presence or absence of H-1PV, at MOIs of 2.5 or 10. After incubation, wells were washed three times with PBS, and 0.5 mL of fresh complete medium (± dox) was added. At each indicated time point, supernatants were collected and diluted 1:2 in PBS in individual 1.5 mL tubes for analysis of released virus. To monitor intracellular viral genome accumulation, cells were seeded in 24-well plates at a density of 2 × 10⁵ cells per well in 0.5 mL complete DMEM ± dox, aiming for a full monolayer by the next day. Plates were prepared with four replicate wells per condition, and a dedicated plate was used for each time point (15 min, 1 h, 2 h, and 4 h). Cells were infected with 250 µL of IM ± dox, ± H-1PV at MOIs of 2.5 or 10. At each time point, supernatants were aspirated, cells were washed four times with PBS, and detached with 250 µL trypsin for 5 minutes at 37°C. The reaction was stopped by adding 250 µL of complete DMEM, and cells were thoroughly dissociated. Four hundred microliters of each cell suspension were diluted in 600 µL PBS (cell fraction) and transferred into 1.5 mL microtubes. Samples (1 mL total) were freeze-thawed three times using liquid nitrogen, then centrifuged at 12,000 × g for 10 minutes at 4°C. Supernatants were collected for qPCR analysis. For quantification of viral genomes, MicroAmp™ Fast Optical 96-Well Reaction Plates (Invitrogen – 4346906) were filled with 18 µL of qPCR mix, composed of 10 µL SsoFast™ EvaGreen® Supermix (Bio-Rad – 1725204), 0.66 µL of each oligonucleotide (10 µM): Forward: 5′-GCGCGGCAGAATTCAAACT-3′ and Reverse: 5′-CCACCTGGTTGAGCCATCA-3′, and 7.88 µL DNAse/RNAse-free water. Then, 2 µL of each sample was added to individual wells. A standard curve was generated using serial dilutions of the pSR19 plasmid, kindly provided by Dr. Jean Rommelaere, which contains the full H-1PV genome. Known copy numbers of pSR19 were used to calculate genome copy numbers in each sample. Fast RT-qPCR was performed using the StepOnePlus™ Real-Time PCR System (Applied Biosystems – 4376600), and data were analyzed with StepOne Software v2.3.

### Immunofluorescence and viral proteins detection

Cells were seeded in 24-well plates at a density of 4.5 × 10⁴ to 8 × 10⁵ cells per well on 14mm coverslips (Knittel Glass - 100041) with 1 mL of complete DMEM, with or without dox, and incubated at 37°C. The following day, cells were incubated for 1 hour at 37°C with 250 µL of infection medium, in the presence or absence of H-1PV at MOIs of 1 or 2.5. After the incubation, the cells were washed three times with PBS and replaced with 1 mL of complete medium (± dox) and incubated for an additional 24 hours at 37°C. At the end of the incubation, cells were washed with PBS and fixed with PBS-PFA 4% (Electron Microscopy Science - 15710) for 15 minutes at room temperature. After fixation, cells were washed three times with PBS, permeabilized with PBS-Triton X-100 0.25% (Sigma-Aldrich - 93443) for 20 minutes at room temperature, washed three times with PBS, and blocked in PBS-BSA 5% (Euromedex - 04-100-812-E) for 30 minutes at room temperature. For the immunodetection of proteins of interest, cells were incubated with primary antibody diluted in 200 µL of PBS-BSA 1% for 2 hours at room temperature with stirring. After primary antibody incubation, cells were washed three times for 5 minutes with PBS and then incubated with secondary antibody diluted in 200 µL of PBS-BSA 1% for 1 hour at room temperature with stirring. Cells were then incubated with PBS containing DAPI at 0.5 µg/mL (Euromedex - 1050A) for 10 minutes at room temperature. After DAPI staining, cells were washed three times for 5 minutes with PBS. Finally, cells were mounted on glass slides with 10 µL of fluorescent mounting medium (Dako - 53023). Image acquisition was performed using a widefield microscope (ZEISS - AxioObserver Z1) with a 10X objective or a confocal microscope (ZEISS - LSM880 Fast Airyscan) with a 63X objective. Images were analyzed with ImageJ 1.53o software. DAPI staining was used to determine the number of cells per field, and fluorescence intensity and area were normalized accordingly. All antibodies used, including dilutions, are listed in Tablexxx. Rabbit antisera directed against H-1PV capsid were kindly provided by Dr. Jürg Nüesch (DKFZ, Heidelberg, Germany).

### Viral particles co-localization studies and binding kinetics

For co-localization studies, cells were seeded in 24-well plates at a density of 4.5 × 10⁴ cells per well on 14 mm coverslips, in 1 mL of complete DMEM supplemented with or without dox, and incubated for 48 hours at 37 °C. To assess viral entry, cells were pre-incubated for 15 minutes on ice, followed by a 1-hour incubation on ice with 250 µL of infection medium containing H-1PV at a MOI of 250, in the presence or absence of dox. Cells were transferred to 37 °C to allow synchronized viral entry. For binding kinetic assays, cells were first incubated for 2 hours at 37 °C with 0.5 mL of complete DMEM, with or without dox, and with or without hydroxy-dynasore, a dynamin inhibitor. Cells were then incubated with 250 µL of IM supplemented with the same conditions (± dox, ± hydroxy-dynasore) and H-1PV at a MOI of 250, and maintained at 37 °C. At various time points ranging from 0 minutes to 4 hours post-infection, cells were extensively washed with PBS to remove unbound virus and immediately processed for immunofluorescence staining, as described previously.

### GFP-Trap

On day 0, cells were seeded in 100 mm dishes at a density of 1.5 × 10⁶ cells per dish in 10 mL of complete DMEM supplemented with doxycycline. On day 1, H-1PV was added directly to the culture medium at a MOI of 2.5. On day 4, protein extraction was performed as previously described. GFP-Trap was then carried out using agarose beads coupled with highly specific anti-GFP nanobodies (Proteintech – gta), following the manufacturer’s instructions. Briefly, cell lysates were diluted with 300 µL of dilution buffer (10 mM Tris-HCl, 150 mM NaCl, 0.5 mM EDTA, pH 7.5) supplemented with 25 µL of GFP-Trap beads, and incubated for 1 hour at 4 °C on a rotating wheel. Beads were collected by centrifugation at 2,500 × g for 2 minutes at 4 °C and washed four times with wash buffer (dilution buffer supplemented with 0.05% NP-40) to remove unbound proteins. Finally, bead-bound proteins were eluted by heating for 5 minutes at 95 °C in 2X Laemmli buffer. For Western blot analysis, 50 µL of input lysate (I), flow-through (FT, unbound proteins), and bead-bound fraction (B) were loaded per sample.

### Statistics and Reproducibility

No sample size calculation was performed. All experiments were conducted with at least three biologically independent replicates, and each experiment included a minimum of three technical replicates. No data were excluded from the analysis. Calculations were performed using Microsoft Excel Professional Plus 2016. Statistical analyses were carried out using GraphPad Prism version 10.1.2. For comparisons involving two or fewer groups, unpaired two-tailed Student’s t-tests were used. For comparisons involving more than two groups, ordinary one-way ANOVA was performed. In both cases, a 95% confidence interval was applied. In micrographs, p-values are indicated as follows: ns for non-significant (p > 0.05), * for p ≤ 0.05, ** for p ≤ 0.01, *** for p ≤ 0.001, and **** for p ≤ 0.0001. Unless otherwise specified, error bars on micrographs represent standard deviation (S.D.).

## Data availability

All data generated or analyzed during this study are included in this published article and its supplementary information files. Access to raw data will be made available upon reasonable request to the corresponding author.

## Results

### CRISPR/Cas9 KO screen identifies host genes for H-1PV infection in PDAC cells

To uncover host factors essential for H-1PV infection, we performed a genome-wide CRISPR-Cas9 knockout screen using the lentiviral GeckoV2 sgRNA library in the PDAC087T primary pancreatic cancer primary culture (Fig. 1A). This model was selected for its biological relevance, robust proliferation, amenability to lentiviral transduction, and susceptibility to H-1PV-mediated oncolysis. Following Cas9 expression and library transduction, cells were either mock-treated or infected with H-1PV. Genomic DNA was extracted one week post-infection, and integrated sgRNA sequences were PCR-amplified and subjected to next-generation sequencing. MAGeCK analysis identified 19 candidate genes whose loss conferred resistance to H-1PV infection (Fig. 1B). To validate these candidates, we conducted a secondary screen in which each gene was targeted using two independent sgRNAs expressed transiently. Three non-targeting sgRNAs served as negative controls. Transfected cells were challenged with H-1PV, and cell viability was quantified 48 hours later (Fig. 1C-D). Knockout of *GAD2*, *MSLN*, *CORO7*, and *ACTL7B* did not significantly alter sensitivity to H-1PV, suggesting these represent false positives or context-dependent hits. In contrast, loss of *SLC35A1*, *SLC35A2*, *CMAS*, *ST3GAL4*, *TGOLN2*, *UNC50*, *GOSR1*, *VPS51*, *VPS54*, *PTAR1*, *COG1*, *COG3*, *COG4*, or *SEC61B* led to reproducible protection from virus-induced cytotoxicity in at least one sgRNA per gene, supporting their functional involvement in H-1PV infection.

**Figure 1:**
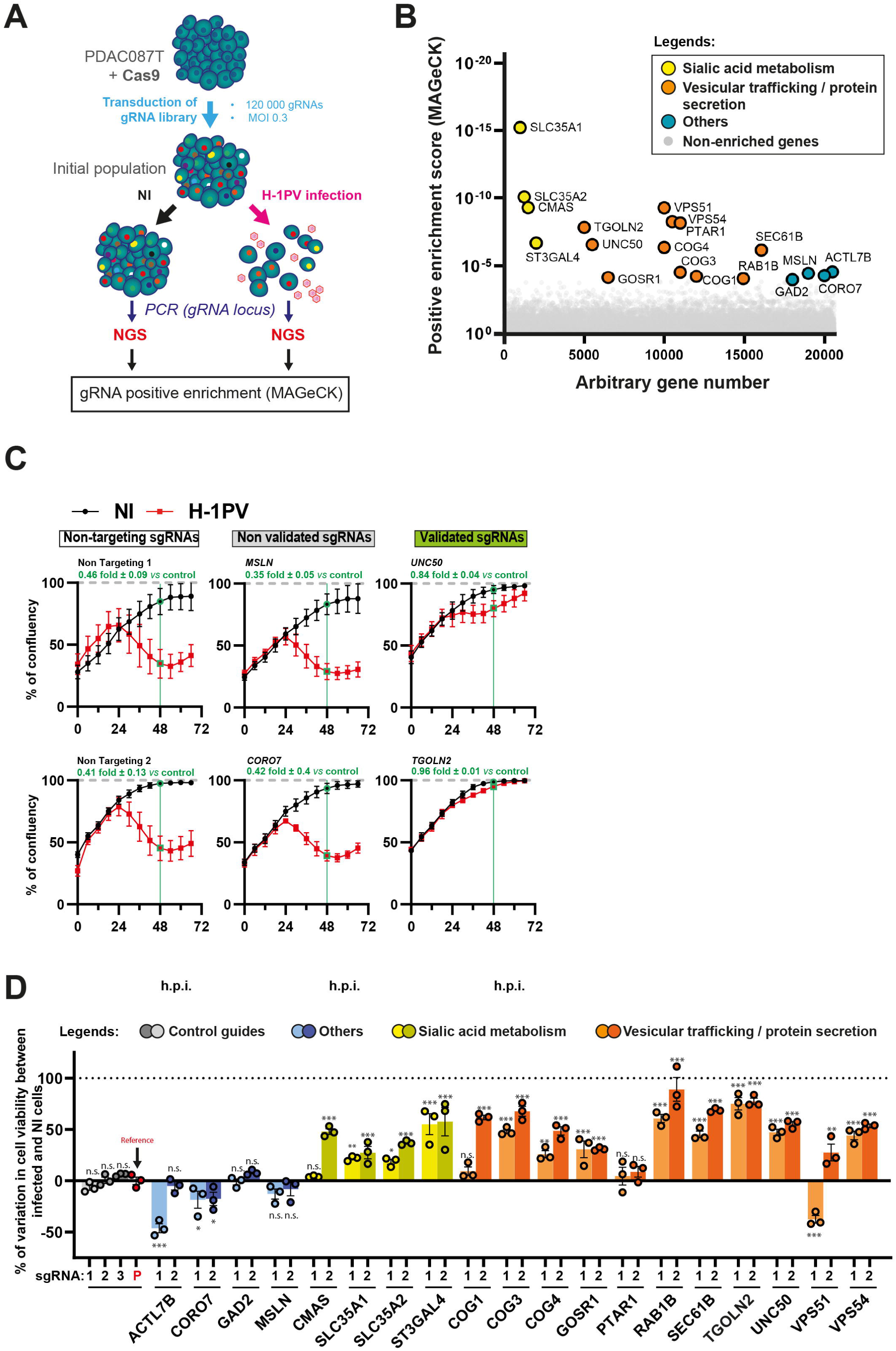
CRISPR/Cas9 KO screen identifies TGOLN2 as a host gene for H-1PV infection. **A.** CRISPR/Cas9 KO screening strategy of H-1PV host genes. PDAC087T cells constitutively expressing Cas9 nuclease enzyme were transduced with the lentiviral pooled sgRNA library GeckoV2 at 0.3 MOI to obtain one gene KO per cell. Cells were then infected with H-1PV or not (NI) at a MOI of 10 and sgRNA content of each population was determined by PCR of sgRNA locus, NGS sequencing and MAGeCK sgRNA enrichment analysis. **B.** Micrograph showing each individual gene screened with an arbitrary gene number in x-axis and the sgRNA positive enrichment score determined with MAGeCK algorithm in y-axis. Genes under positive enrichment score cut-off are drawn with gray dots; genes above cut-off were annotated and grouped by family: sialic acid metabolism (yellow dots), vesicular trafficking / protein secretion (orange dots) and others (teal dots). **C.** Representative validation of gRNA screening results. Examples are shown for control non-targeting gRNAs, for non-validated gRNAs that did not affect viral cytotoxicity (targeting *MSLN*, *CORO7*), and for validated gRNAs that abolished viral lytic activity (targeting *UNC50*, *TGOLN2*). An MOI of 5 was used for H-1PV infection. Cell confluence was monitored using the IncuCyte live-cell imaging system. Data represent the mean ± S.D. of three independent biological experiments performed in triplicate. **D.** Individual validation of genes above positive enrichment score threshold. PDAC087T-Cas9 cells were transduced to express stably individual sgRNAs of each gene to validate (2 sgRNA per gene and 3 non-targeting human control sgRNAs), then cells were infected or not with H-1PV at MOI 5. 48h post infection, cell viability assay was performed and survival ratio (infected / NI) calculated for each sgRNA. Mean ratios of the three non-targeting human controls sgRNA were pooled (P, red dots) and use as reference to calculate resistance to H-1PV conferred by each individually tested sgRNA (% variation of infected / NI). Negative value stands for increased sensitivity to infection. Dots and bars colors correspond to target gene family: non-targeting (gray and light gray), sialic acid metabolism (yellow), vesicular trafficking (orange) and others (blue). For statistical analysis, Fisher’s LSD multiple comparisons test was performed.

To gain insight into the functional landscape of candidate H-1PV host factors, we conducted gene ontology enrichment analysis using the PANTHER classification system (Table 5). This analysis highlighted several significantly enriched biological processes, including nucleotide sugar transport, retrograde trafficking at the trans-Golgi network (TGN), intra-Golgi vesicular transport, sialic acid metabolism, and coatomer protein complex I (COPI)-mediated anterograde transport from the endoplasmic reticulum (ER) to the Golgi apparatus. Notably, many identified genes are involved in vesicle-mediated trafficking, suggesting a model in which H-1PV co-opts host transport machinery for efficient intracellular trafficking and infection. Among the candidate genes, we focused on *TGOLN2*, which encodes the glycoprotein TGN46, a conserved sialylated glycoprotein predominantly localized to the TGN, where it functions in the sorting of cargo destined for secretion^20^. Furthermore, a fraction of TGN46 also traffics to the plasma membrane, where it interfaces with components of clathrin-mediated endocytosis (CME)^27^, a process that was shown to be critical for efficient H-1PV entry^17^. Given the strong protective phenotype observed upon *TGOLN2* knockout (Fig. 1C-D) and its known roles in endocytic and secretory pathways, we hypothesized that TGN46 facilitates a key step in H-1PV entry or intracellular trafficking, warranting further mechanistic investigation.

### *TGOLN2* knockout and overexpression models demonstrate a strong dependency of H-1PV on TGN46 for successful infection

To determine whether TGN46 is required for H-1PV infection, we first performed loss-of-function experiments using gene knockout (KO). We generated PDAC087T cells deficient for *TGOLN2* (PDAC087T-KO) using a non-integrative strategy to minimize permanent genetic alterations and reduce off-target effects associated with constitutive Cas9 expression^28^ (Fig. S1A). Efficient TGN46 depletion was confirmed at the protein level by western blotting (Fig. S1B) and immunofluorescence (Fig. S1C). PDAC087T-KO cells demonstrated complete protection from infection at MOI 2.5 or 10 (Fig. 2A), whereas control cell numbers decreased by 35% ± 9% (p < 0.004) and 67% ± 9% (p < 0.001), respectively (Fig. 2A). We next extended this analysis to HeLa cervical cancer cells, a model known to be permissive to H-1PV infection^29^. Western blot confirmed the *TGOLN2* KO (Fig. S1D). Viability assays performed 48 hours post-infection showed that *TGOLN2* KO resulted in a significant reduction in H-1PV-induced oncolysis compared with their respective control cells (Fig. S1E), indicating that TGN46 may function as a conserved host factor for H-1PV infection. We then measured viral genome content in wt and PDAC087T-KO cells following infection with H-1PV. Consistent with the reduction in oncolysis, viral replication was strongly impaired in the absence of TGN46. Quantification of viral DNA by qPCR revealed a significant decrease in intracellular H-1PV genome copies in *TGOLN2*-deficient cells compared with controls (−94% ± 0.5%, p < 0.01 and −82% ± 5.6%, p < 0.001 for MOI 2.5 and MOI 10, respectively; Fig. 2B). To assess whether this defect translated into altered production of infectious viral particles, we next quantified extracellular H-1PV virions in culture supernatants. TGN46-deficient PDAC087T cells released significantly fewer viral particles compared with control cells, as measured by qPCR-based particle quantification (−73% ± 31%, p < 0.001, and −71% ± 11%, p < 0.001, at MOI 2.5 and MOI 10, respectively Fig. 2C). These findings indicate that TGN46 loss impairs intracellular viral genome amplification and limits productive virion generation. We next assessed viral protein expression by immunofluorescence. In control cells, robust expression of NS1 and capsid viral protein was readily detected, with a characteristic intracellular distribution (Fig. 2D-E). In contrast, *TGOLN2*-deficient cells consistently displayed a marked reduction in the proportion of infected cells (−74% ± 18%, p < 0.001 for NS1 and −85% ± 11%, p < 0.001 for capsid proteins, respectively Fig. S1F-G), consistent with impaired viral gene expression. We then assessed whether TGN46 overexpression was sufficient to modulate viral infection and efficacy. To this end, we established PDAC087T cells in which TGN46 overexpression is induced upon doxycycline (dox) treatment (Fig. S1H). Western blotting (Fig. S1I) and immunofluorescence (Fig. S1J) for TGN46 validated both cellular models. We next assessed the impact of TGN46 overexpression following infection with H-1PV at various MOI. Proliferation assays indicated that doxycycline treatment did not affect cell growth, and revealed that TGN46 overexpression resulted in 25% ± 4% (p < 0.004) and 31% ± 4% (p < 0.03) reduced viability at MOI 2.5 and 10, respectively (Fig. 2F). Furthermore, intracellular viral genome content was significantly higher in cells overexpressing TGN46, compared with control cells, with a 2.4 ± 0.2-fold (p < 0.01) and 1.5 ± 0.3-fold (p < 0.002) increase at MOI 2.5 and 10, respectively (Fig. 2G). In contrast, viral genome levels detected in the culture supernatant, reflecting released progeny virions, were not significantly altered (Fig. 2H). Together, these findings are consistent with a role for TGN46 in early stages of the H-1PV infectious cycle rather than in late steps of virion production or release. Finally, Immunofluorescence analysis further showed enhanced viral protein expression upon TGN46 overexpression. Twenty-four hours post-infection at MOI 2.5, TGN46-overexpressing cells exhibited a significant increase in the percentage of NS1 (1.7-fold ± 0.5, p<0.05, Fig. 2I and Fig. S1K) and conformed viral capsid-positive cells (3.6-fold ± 0.9, p<0.01, Fig. 2J and Fig. S1L) compared with controls, consistent with increased viral gene expression and productive infection. Taken together, these complementary loss-and gain-of-function experiments identify *TGOLN2*/TGN46 as a host factor that promotes productive H-1PV infection, consistent with a role at an early stage of the viral life cycle, potentially at or upstream of viral entry.

**Figure 2:**
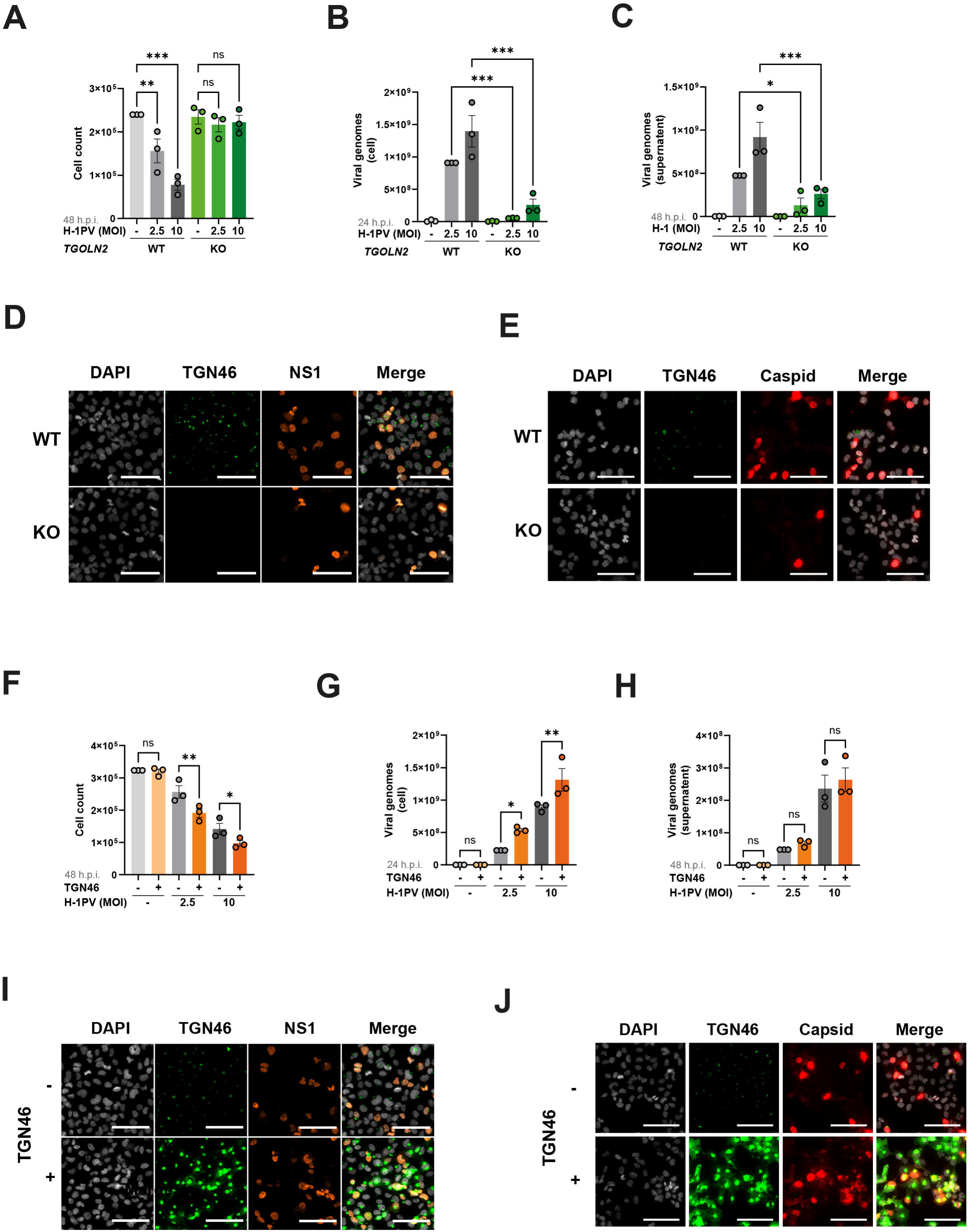
H-1PV rely on TGN46 expression for successful infection and oncolytic activity. (**A, F**) cell counts, (**B, G**) intracellular H-1PV viral genomes content (qPCR targeting the NS1 genomic locus) and (**C, H**) H-1PV viral genomes in the supernatant of PDAC087T-WT and *TGOLN2*-KO cells (**A-C**), and control or of TGN46-overexpressing PDAC087T cells (**F-H**), measured 48h post H-1PV infection. Infections were performed at MOI 2.5 and 10. Statistical comparisons were performed using a two-way ANOVA. (D, E) Immunofluorescence (IF) analysis of PDAC087T-WT and *TGOLN2*-KO cells, and (I, J) control or TGN46-overexpressing PDAC087T cells, 24 h post–H-1PV infection at an MOI of 2.5. Cells were stained with anti-TGN46 antibody (green) and DAPI for DNA counterstaining (light gray). Anti-NS1 antibody (orange) and anti-H-1PV conformational capsid antibody (red) were used to detect viral proteins. Images were acquired using a wide-field microscope with a 20X objective (scale bars: 100 µm).

### TGN46 regulates H-1PV entry and co-localizes with H-1PV virions at the cell membrane

Given TGN46’s involvement in protein secretion and vesicular recycling, we sought to determine whether its role in H-1PV infection might be indirect, affecting endocytosis pathways, or if TGN46 directly contributes to the virus’s attachment or entry into cells. To investigate this, we performed immunofluorescence analyses to assess the steady-state levels of major endocytosis markers of CME in both PDAC087T-KO and PDAC087T cells overexpressing TGN46. Confocal microscopy images were analysed to quantify expression levels, vesicle numbers, and average vesicle sizes using Clathrin Heavy Chain (CHC), Rab5a (early endosome marker), EEA1 (early-intermediate endosome marker), and LAMP1 (lysosomal marker). Single-cell quantification revealed that *TGOLN2* knockout or overexpression did not generally alter the expression levels of these endocytic markers, with the exception of EEA1, whose relative signal was increased in *TGOLN2*-depleted cells (Fig. S2A-B). *TGOLN2* downregulation also induced significant changes in both the number (CHC, Rab5a, EEA1) and size (CHC, Rab5a, EEA1) of endocytic vesicles, whereas TGN46 overexpression had no detectable effect. To assess whether these morphological alterations affected CME functionality, we performed transferrin uptake assays using Alexa-647-conjugated transferrin (TFn-647), which is internalized exclusively via CME^30^. After a 5-minute incubation, no differences in total TFn-647 uptake were observed between *TGOLN2* knockout, TGN46-overexpressing, and wild-type cells (Fig. S2A-B). However, the number of transferrin-positive vesicles was reduced in *TGOLN2*-depleted cells. Together, these data indicate that modulation of TGN46 expression does not impair the overall capacity for CME in PDAC087T cells.

Based on these observations, we next investigated whether TGN46 plays a specific role during the early stages of H-1PV entry. To address this, we performed an H-1PV internalization assay. PDAC087T cells were incubated with H-1PV, and at the indicated time points, cells were washed to remove unbound virus (Fig. 3A). Viral genome content was then quantified by qPCR. At 2 hours post-infection, *TGOLN2*-KO cells contained 45% fewer viral genomes than wild-type controls (Fig. 3A, p<0.005), whereas TGN46-overexpressing cells exhibited more than a twofold increase in viral genome levels as early as 15 minutes post-infection (Fig. 3B). These results indicate that TGN46 enhances early H-1PV entry in a dose-dependent manner. To further investigate the role of TGN46 during H-1PV entry, we adapted a previously described synchronization protocol^17^ to visualize the interaction between TGN46 and H-1PV using an antibody recognizing conformational capsid epitopes in intact virions. Vesicular trafficking was transiently inhibited in PDAC087T cells by incubation at 4°C for 15 minutes, followed by exposure to a high viral input (MOI 250) for 1 hour at 4°C to allow virus binding without internalization. Viral entry was subsequently initiated by shifting cells to 37°C, and cells were fixed at 5 and 20 minutes post-release. TGN46 and H-1PV capsid were detected by immunostaining, and samples were imaged using super-resolution confocal microscopy. In PDAC087T cells overexpressing TGN46, a pronounced accumulation of H-1PV capsid was observed at the cell periphery as early as 5 minutes post-release (Fig. 3C, left panel). At 20 minutes post-release, extensive viral internalization was detected in TGN46-overexpressing cells, with significant co-localization of the viral capsid within TGN46-positive vesicles, consistent with trafficking through TGN-associated compartments (Fig. 3C, right panel). Comparable capsid-TGN46 co-localization was also observed in wild-type PDAC087T cells, although it occurred at later time points (60 minutes post-release) and with reduced TGN46 accumulation (Fig. S2C). Notably, cells in prophase, a stage characterized by fragmentation of the TGN and disruption of vesicular trafficking, displayed an accumulation of TGN46/capsid-positive vesicles (Fig. S2D), consistent with a block at an early stage of viral entry.

**Figure 3:**
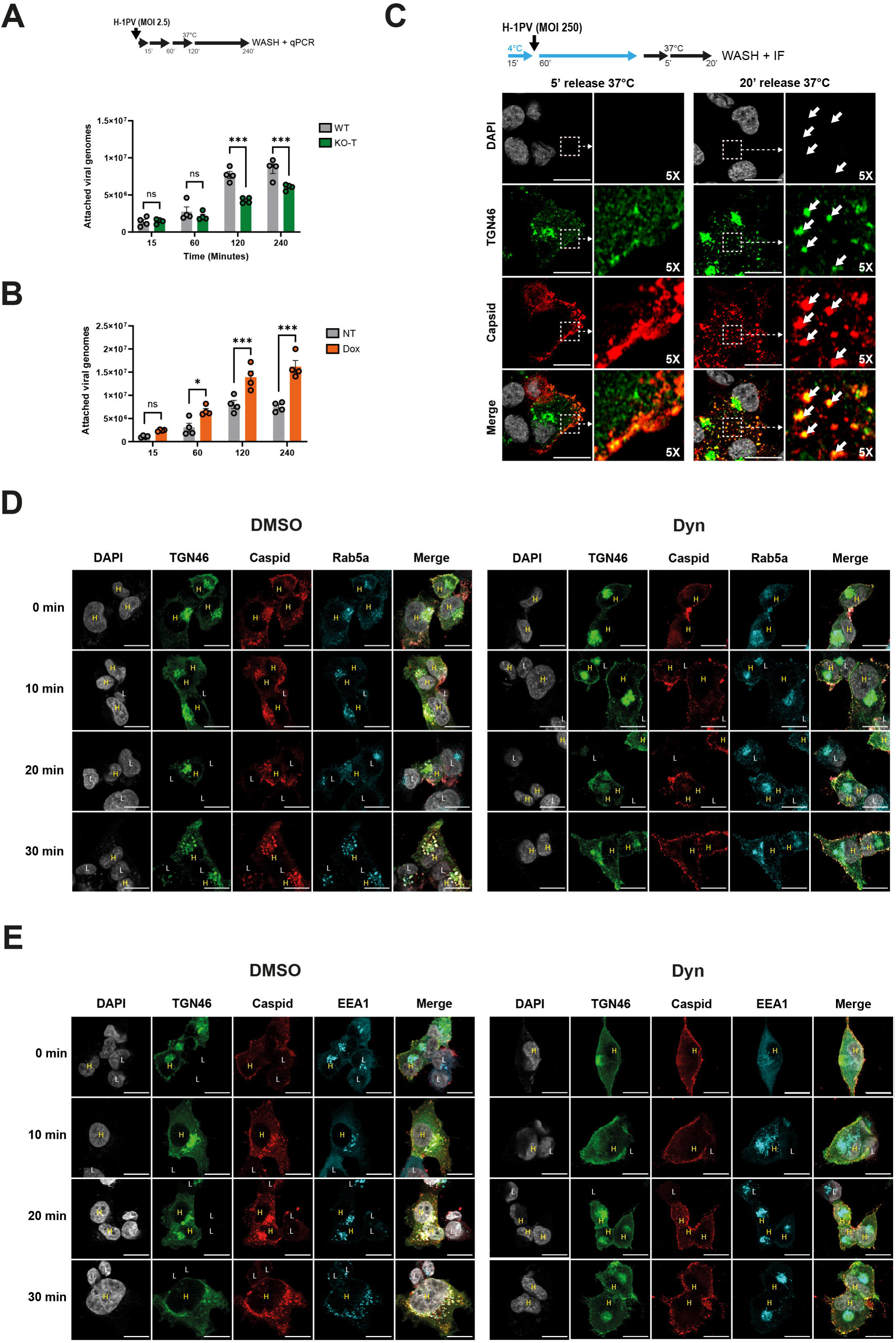
Modifications in TGN46 expression level do not impact CME but controls H-1PV entry. PDAC087T-WT (WT) cells, or PDAC087T cells KO for *TGOLN2* (KO-T, **A**), and PDAC087T cells overexpressing (+) or not (-) TGN46 (**B**) were infected with H-1PV at MOI 2.5 and incubated 15, 60, 120 or 240 min at 37°C followed with extensive wash with PBS, to remove unbound virus, and internalized viral genomes were quantified by qPCR against NS1 genomic locus. Statistical comparisons were performed using a two-way ANOVA**. C.** Cells were pre-incubated for 15 min at 4°C and then infected with H-1PV at an MOI of 250 for 1 h at 4°C to allow viral binding to the cell surface. Cells were subsequently shifted to 37°C for 5 or 20 min, extensively washed with PBS, fixed, and processed for IF staining of TGN46 (green), viral capsid (red), and DAPI for DNA counterstaining (light gray). Images were acquired using a confocal microscope with a 63× objective (scale bars, 20 µm). Regions of interest were magnified 5-fold. PDAC087T-*TGOLN2*-KO cells reconstituted with a doxycycline-inducible TGN46-GFP construct and pretreated for 30 min with DMSO (control, **D** and **E**, left panels) or 100µM hydroxy-dynasore (Dyn, **D** and **E**, right panels). Cells were infected with H-1PV at an MOI of 250 for the indicated times, and stained with antibodies against TGN46 and viral capsids (D, E), together with antibodies against Rab5a (**D**) or EEA1 (**E**) and DAPI for DNA counterstaining (light gray). Images were acquired using a confocal microscope with a 63× objective (scale bars, 20 µm).

To determine whether TGN46 interacts with the H-1PV virions at the cell surface prior to internalization, as would be expected for a functional viral receptor, we explored experimental conditions that stabilize TGN46 at the plasma membrane. Owing to its rapid recycling kinetics, visualization of surface-localized TGN46 under steady-state conditions is technically challenging. Previous studies have shown that inhibition of dynamin I impairs the scission of clathrin-coated pits, resulting in the accumulation of TGN46 at the cell surface and a concomitant block in CME^31^. We therefore treated PDAC087T cells with increasing concentrations of hydroxy-dynasore, a reversible, non-competitive inhibitor of dynamin I, to phenocopy this condition. To validate CME inhibition, we monitored uptake of transferrin-Alexa647 (TFn-647) after 5 minutes of incubation and quantified fluorescent intensity and area by immunofluorescence (Fig. S2E-F). As expected, TFn-647 internalization was dose-dependently inhibited by dynasore, with near-complete blockade at 100 µM (−95% ± 1.5%, p<0.001, Fig. S2F). In parallel, total TGN46 levels remained unchanged (Fig. S2E and S2G), whereas the spatial distribution of the protein shifted, with a significant increase in membrane-associated signal, consistent with plasma membrane accumulation (3.1-fold ± 0.2, p<0.001, Fig. S2E and S2H). We next assessed whether surface-localized TGN46 engages the H-1PV capsid. PDAC087T-*TGOLN2*-KO cells reconstituted with a doxycycline-inducible *TGOLN2*-GFP construct were used to compare TGN46-high and TGN46-low cells. Cells were pretreated with dynasore or DMSO and exposed to H-1PV under synchronized infection conditions. At defined time points, cells were immunoassayed for H-1PV capsid proteins and either Rab5a (Fig. 3D) or EEA1 (Fig. 3E) to label early endosomes; TGN46-GFP was detected directly via GFP fluorescence. Under control conditions, TGN46-high cells showed greater accumulation of H-1PV capsids compared to TGN46-low cells, with both TGN46 and capsid signals progressively localizing to Rab5a+ and EEA1+ endosomes. In contrast, dynasore treatment prevented internalization, resulting in co-localization of TGN46 and capsids at the plasma membrane, without detectable overlap with Rab5a or EEA1 (Fig. S2I). These results indicate that TGN46 engages H-1PV at the cell surface within Rab5a-enriched microdomains prior to CME. Collectively, our data identify TGN46 as a novel entry factor for H-1PV that transiently localizes to the plasma membrane to mediate viral engagement before internalization.

### TGN46 domain I interacts with H-1PV capsid proteins for successful infection

To elucidate the mechanism underlying the interaction between TGN46 and H-1PV, we used AlphaFold 2.0 to predict potential structural interfaces between the lumenal domain of TGN46 and the viral capsid proteins. We submitted the full lumenal sequence of TGN46, which behaves as an intrinsically disordered region (IDR) and is therefore challenging to model, together with the full VP2 sequence, to AlphaFold (Fig. 4A). This analysis generated five models, which were evaluated using the predicted local distance difference test (pLDDT), predicted aligned error (PAE), predicted template modeling score (pTM), and interface pTM (ipTM) (Fig. 4B). As expected, the high flexibility of the IDR resulted in low ipTM values; however, the top-ranked model showed moderate confidence (pLDDT = 62.7, pTM = 0.618, ipTM = 0.212). Notably, all five models converged on a predicted interaction between domain I of TGN46 and the sialic acid (SIA)-binding pocket of VP2 (Fig. 4B). To improve modelling accuracy, we truncated the input sequences to include only the first 50 amino acids of TGN46 domain I and the SIA-binding pocket of VP2. This refinement resulted in a marked increase in prediction scores (pLDDT = 87.6, pTM = 0.855, ipTM = 0.372), supporting a higher-confidence interaction model between TGN46 domain I and VP2.

**Figure 4:**
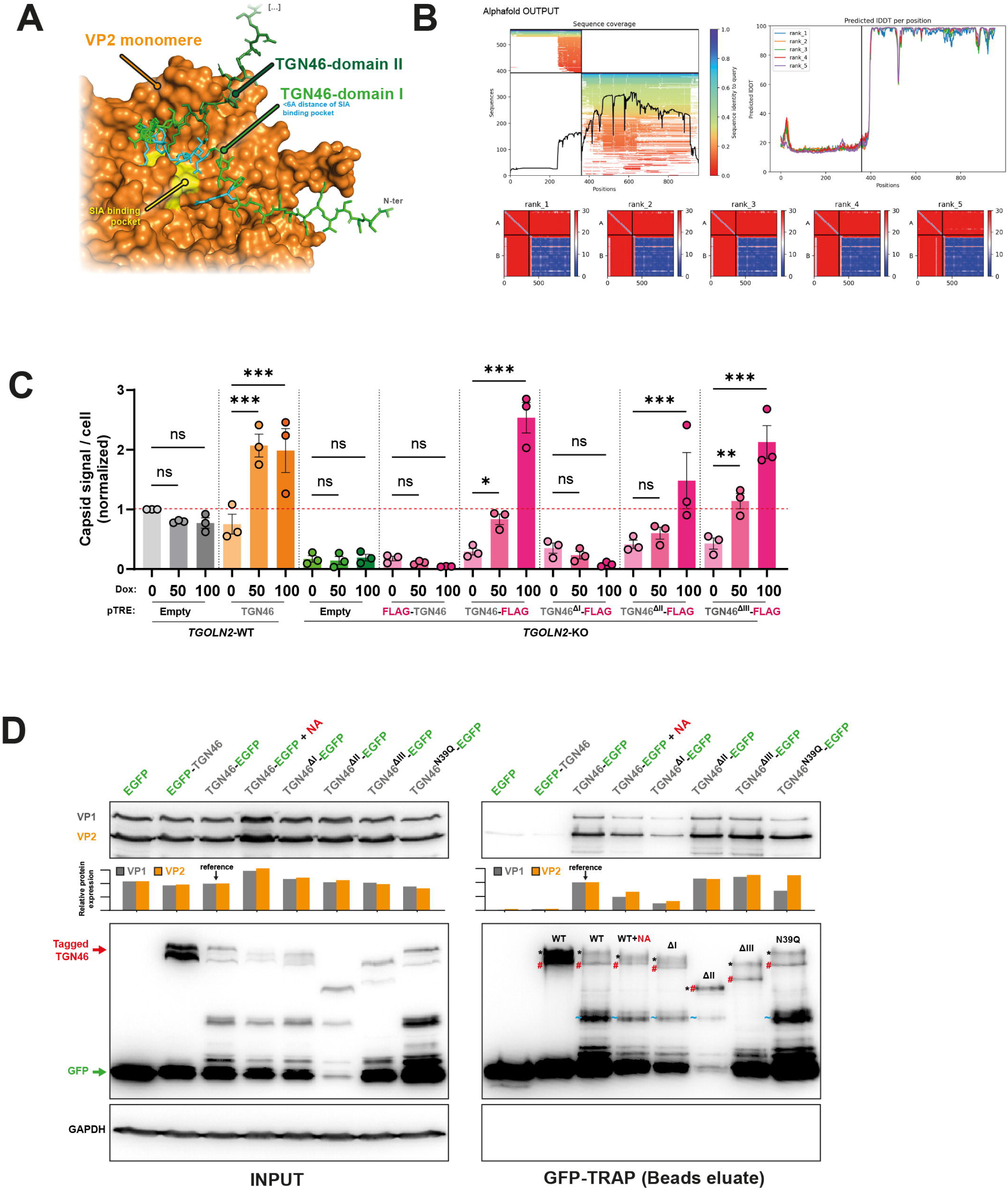
H-1PV infection requires interaction of capsid proteins with sialic acid motifs present in TGN46 N-terminal domain I. **A**. Predicted model of the interaction between the full luminal domain of TGN46 and H-1PV capsid proteins generated using AlphaFold 2.0. **B**. Model evaluation based on predicted local distance difference test (pLDDT), predicted aligned error (PAE), predicted template modeling score (pTM), and interface predicted template modeling score (ipTM). **C.** PDAC087T-WT cells (gray bars); PDAC087T cells with dox-inducible TGN46 expression (orange bars), PDAC087T-*TGOLN2*-KO cells (green bars); and PDAC087T-*TGOLN2*-KO cells co-expressing dox-inducible FLAG-TGN46, TGN46-FLAG, TGN46ΔI-FLAG, TGN46ΔII-FLAG, or TGN46ΔIII-FLAG (pink bars) were treated with 0, 50, or 500 µg/mL dox) and infected with H-1PV at an MOI of 2.5 for 24 h. Cells were fixed and stained with an antibody against H-1PV conformational capsid proteins and DAPI for DNA counterstaining (light gray). Images were acquired using a confocal microscope with a 20× objective. Total capsid fluorescence intensity per cell was quantified and normalized to the control condition (red dotted line). Data represent mean ± SD of three independent experiments performed in triplicate. Statistical analysis was performed using uncorrected Fisher’s LSD test**. D.** PDAC087T-*TGOLN2*-KO cells transduced with lentiviral vectors encoding EGFP, EGFP-TGN46, TGN46-EGFP, TGN46ΔI-EGFP, TGN46ΔII-EGFP, TGN46ΔIII-EGFP, or TGN46N39Q-EGFP were infected with H-1PV at an MOI of 2.5. In addition, PDAC087T-*TGOLN2*-KO cells expressing TGN46-EGFP were treated with neuraminidase prior to infection. Two days post-infection, cellular proteins were extracted and incubated with GFP-Trap beads. Bead-bound proteins were resolved by western blot. Input whole-cell lysates and GFP-Trap co-immunoprecipitated proteins were analyzed by western blot for EGFP, VP1, and VP2.

To functionally validate these predictions, we conducted TGN46 rescue experiments in PDAC087T *TGOLN2* KO cells. Cells were transduced with lentiviral vectors encoding dox-inducible *TGOLN2* fused to a FLAG tag at either the N- or C-terminus (Fig. S3A). Control conditions included empty vectors or EGFP alone. Truncated versions of TGN46 lacking individual lumenal domains (ΔI, ΔII, ΔIII) were also generated, with the signal peptide preserved. Immunofluorescence confirmed proper expression of all constructs (Fig. S3B). We then assessed the ability of FLAG-tagged TGN46 variants to rescue H-1PV infection in PDAC087T-KO cells. Following H-1PV infection, infectivity was measured by immunofluorescence detection of conformational capsid proteins. C-terminally FLAG-tagged TGN46 fully restored H-1PV infectivity in PDAC087T-KO cells (Fig. 4C). We also generated a C-terminally GFP-tagged version of TGN46, which was validated for proper expression (Fig. S3C) and similarly rescued infection in PDAC087T-KO cells (Fig. S3D). In contrast, N-terminally tagged constructs failed to restore infectivity (Fig. 4C, Fig. S34). Among the domain-deletion mutants, the ΔII and ΔIII variants supported infection at levels comparable to full-length TGN46, whereas the ΔI mutant failed to restore infectivity, indicating that luminal domain I (32 amino acids) is essential for H-1PV infection (Fig. 4C).

To assess whether TGN46 physically associates with the H-1PV capsid, we performed GFP-TRAP co-immunoprecipitation assays in PDAC087T-WT cells transduced with EGFP-tagged TGN46 constructs. Following H-1PV infection, GFP fusion proteins were immunoprecipitated using anti-GFP nanobodies (Fig. S3E). Western blot analysis revealed that the viral capsid proteins VP1 and VP2 co-immunoprecipitated specifically with C-terminally tagged TGN46 variants, indicating that TGN46 and the H-1PV capsid belong to the same molecular complex (Fig. 4D). Notably, under these conditions, TGN46 did not associate with the viral protein NS1 nor with galectin-1 (Fig. S3H). Domain-deletion analysis showed that TGN46 variants lacking domains II or III retained capsid interaction, whereas deletion of domain I (ΔI) reduced VP1 and VP2 binding by 75% and 65%, respectively (Fig. 4D). Considering the critical role of protein glycosylation in H-1PV infection^14^, we investigated the contribution of post-translational modifications in TGN46 and H-1PV interaction. TGN46-EGFP expressing cells were treated with neuraminidase (NA), that cleaves sialic acid residues from glycoproteins and glycolipids on the cell surface. We found that NA treatment significantly inhibited H-1PV entry (−53% ± 15 %, p<0.05, Fig. S3I) and induced a mobility shift of mature TGN46 (Fig. 4D). Importantly, NA treatment decreased co-immunoprecipitation of VP1 and VP2 by 50% and 35%, respectively (Fig. 4D). To further probe the role of glycosylation in TGN46-mediated H-1PV entry, we mutated the sole predicted N-glycosylation site within domain I by at asparagine 39 (N39) to glutamine (Q), a conservative substitution that prevents glycosylation while preserving local structure. This N39Q mutation reduced VP1 binding by 35% without affecting VP2 interaction, indicating that this glycosylation contributes to the VP1-TGN46 association (Fig. 4D). Together, these results demonstrate that TGN46 binds H-1PV capsid proteins through domain I, and confirms that sialylated glycosylation is an important determinant of this interaction.

### TGN46 is a key determinant of H-1PV entry and oncolytic activity in PDAC cells

Given that mechanistic studies have collectively identified TGN46 as a critical mediator of H-1PV entry, we next examined whether this dependency dictates the specificity and the oncolytic efficacy of H-1PV in PDAC cells. H-1PV entry was quantified across a panel of PDAC-derived cell lines following infection with an MOI of 200 of the virus, using normal pancreatic ductal epithelial (HPDE) cells as a non-malignant control. HPDE cells exhibited remarkably low levels of viral entry (1.3×10^4^ a.u. ± 0.7×10^4^), which were significantly lower than those observed in Capan-2 (4×10^4^ a.u. ± 2.7×10^4^, p < 0.05), BxPC-3 (9×10^4^ a.u. ± 2.4×10^4^, p < 0.0001), Mia PaCa-2 (1.1×10^5^ a.u. ± 2.3×10^4^, p < 0.0001), and Panc-1 cells (2.2×10^5^ a.u. ± 4.5×10^4^, p < 0.0001), the latter showing the highest susceptibility to infection (p<0.0001, Fig. 5A). In contrast, AsPC-1 cells did not show a significant difference in H-1PV entry compared to HPDE cells (2.5×10^4^ a.u. ± 0.6×10^4^, p = 0.4; Fig. 5A). We then investigated the half-maximal inhibitory concentration (IC₅₀) of H-1PV on the same panel of cells. Normal HPDE exhibited pronounced resistance to H-1PV, with no measurable IC₅₀ (Fig. 5B). Among PDAC cells, AsPC-1 (IC₅₀ MOI = 193.2 ± 2.3) and Capan-2 (IC₅₀ MOI = 132.6 ±2.1) were classified as resistant, while BxPC-3 (IC₅₀ MOI = 19.6 ±1.3), Mia PaCa-2 (IC₅₀ MOI = 5.4 ±0.7), and Panc-1 (IC₅₀ MOI = 2.8 ±0.45) displayed marked susceptibility to viral cytotoxicity. These same cell lines were subsequently analysed for the protein expression of TGN46, along with two known modulators of H-1PV entry, galectin-1^15,32^ and laminin subunit gamma-1 (LAMC1)^33^, to explore potential correlations with viral sensitivity. Interestingly, both galectin-1 and LAMC1 were detected in HPDE cells despite their resistance to infection (Fig. S4A). While galectin-1 expression alone showed a positive correlation with H-1PV sensitivity across the panel, LAMC1 levels did not reflect IC₅₀ values (Fig. S4A). Remarkably, TGN46 expression was absent in HPDE cells and positively correlated with H-1PV sensitivity in PDAC-derived cells (Fig. S4A).

**Figure 5:**
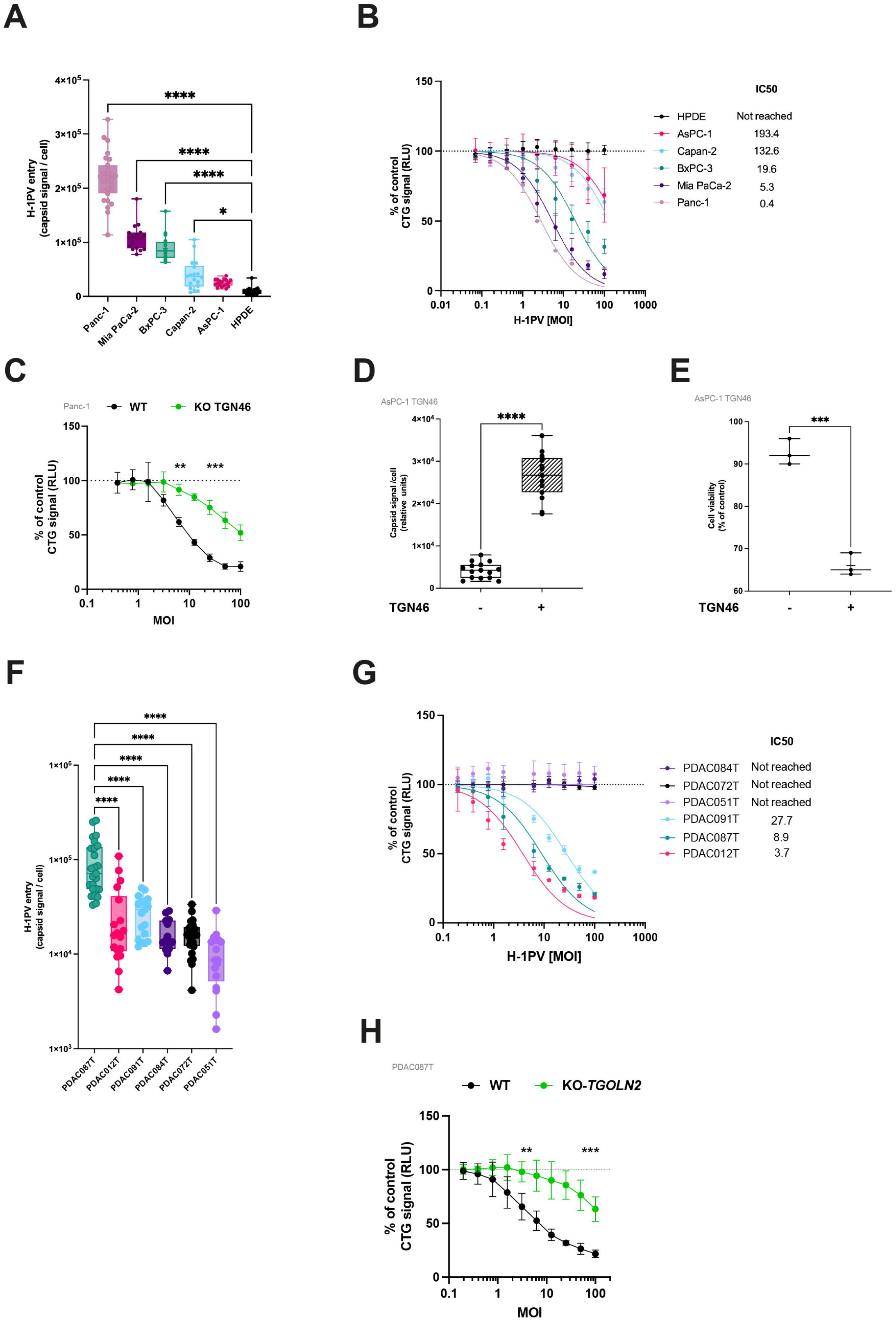
TGN46 drives H-1PV entry and oncolytic efficacy in pancreatic ductal adenocarcinoma cells. **A**. PDAC cell lines were infected with H-1PV (MOI 200) for 2 hours. Cells were then fixed and stained with an antibody targeting the conformational H-1PV capsid protein. Normal pancreatic ductal epithelial (HPDE) cells were used as control. Images were acquired using a confocal microscope (63× objective), and fluorescence intensity was quantified. Data represent the mean ± SD of three independent experiments, each performed in triplicate. Statistical comparisons were performed using a two-way ANOVA. **B**. PDAC cell lines were infected with H-1PV at the indicated MOIs. After 48 hours, cell viability was measured using the CellTiter-Glo (CTG) assay, and IC₅₀ values were calculated. Data represent the mean ± SD of three independent experiments, each performed in triplicate. **C**. Panc-1 cells with *TGOLN2* knockout were generated using CRISPR/Cas9 and infected with H-1PV at the indicated MOIs. After 48 hours, cell viability was measured using CTG assay. Data represent the mean ± SD of three independent experiments, each performed in triplicate. Statistical comparisons were performed using an unpaired t-test. Lentiviral vectors encoding for TGN46 were used to generate AsPC-1 TGN46 cells. Proteins expression was induced or not by doxycycline and cells infected with H-1PV at an MOI of 200 for 2 h (**D**). Cells were then fixed and stained with an antibody targeting the conformational H-1PV capsid protein. Images were acquired using a confocal microscope (63× objective), and fluorescence intensity was quantified. Alternatively, AsPC-1 cells expressing or not TGN46 were infected with H-1PV at an MOI of 50. **E.** After 48 hours, cell viability was measured using the CTG assay. Data represent the mean ± SD of three independent experiments, each performed in triplicate. Statistical comparisons were performed using an unpaired t test. **F.** PDAC primary cells were infected with H-1PV (MOI 200) for 2 hours. Cells were then fixed and stained with an antibody targeting the conformational H-1PV capsid protein. Images were acquired using a confocal microscope (63× objective), and fluorescence intensity was quantified. Data represent the mean ± SD of three independent experiments, each performed in triplicate. Statistical comparisons were performed using a two-way ANOVA. **G**. PDAC primary cells were infected with H-1PV at the indicated MOIs. After 48 hours, cell viability was measured using the CTG assay, and IC₅₀ values were calculated. Data represent the mean ± SD of three independent experiments, each performed in triplicate. **H**. PDAC087T primary cells with *TGOLN2* knockout were generated using CRISPR/Cas9 and infected with H-1PV at the indicated MOIs. After 48 hours, cell viability was measured using CTG assay. Data represent the mean ± SD of three independent experiments, each performed in triplicate. Statistical comparisons were performed using an unpaired t-test.

To further investigate the contribution of TGN46 to H-1PV infectivity, we deleted *TGOLN2* in highly susceptible, TGN46-positive Panc-1 cells using CRISPR/Cas9. Loss of TGN46 expression was confirmed by Western blot (Fig. S4B) and resulted in a 7.5-fold increase in the H-1PV IC₅₀ (p < 0.0001; Fig. 5C), indicating reduced viral sensitivity. In mirror studies, we conditionally overexpressed TGN46 in TGN46-low, H-1PV resistant AsPC-1 cells. Dox addition successfully induced TGN46 expression (Fig. S4C, p<0.0001), and resulted in an a 6.2-fold induction of H-1PV entry (Fig. 5D, p<0.0001) and significant sensitization to H-1PV infection (+35% ± 8, p<0.01, Fig. 5E). We extended our analysis to primary PDAC-derived cultures, as these may better capture tumour heterogeneity than established cell lines and enhance clinical relevance. We quantified H-1PV entry and found that PDAC087T cells exhibited high levels of viral entry (9.7 × 10⁴ a.u. ± 1.3 × 10⁴, p < 0.0001; Fig. 5F). Interestingly, these cells expressed the highest protein levels of TGN46 and galectin-1 (Fig. S4D). We then measured the IC₅₀ of H-1PV on primary PDAC cells. IC_50_ was not reached for PDAC084T, PDAC072T and PDAC051T cultures, that were classified as resistant, while PDAC091T (IC₅₀ MOI = 27.7 ±1.3), PDAC087T (IC₅₀ MOI = 8.9 ±0.7), and PDAC012T (IC₅₀ MOI = 3.7 ±0.45) were highly susceptible to H-1PV-induced cytotoxic effects. We then applied the same strategy used in established cell lines to assess the role of TGN46 in H-1PV infection of primary cells by deleting *TGOLN2* using CRISPR–Cas9 in susceptible PDAC087T cells. Targeting TGN46 expression resulted in a 19-fold increase in the H-1PV IC₅₀ (p<0.0001, Fig. 5H), corresponding to reduced sensitivity to viral infection. Importantly, these results extend our findings from cell lines to primary PDAC cultures and demonstrate that TGN46 contributes not only to H-1PV infection but also to its oncolytic activity. Together, these data warrant further exploration of TGN46 function in experimental models *in vivo*.

### TGN46 is required for H-1PV antitumoral activity *in vivo*

To assess the contribution of TGN46 to H-1PV oncolytic activity *in vivo*, we generated subcutaneous tumors from primary PDAC087T cells in which *TGOLN2* was deleted using CRISPR-Cas9 (Fig. 6A). Efficient and sustained depletion of *TGOLN2* in tumour tissues was confirmed at the protein level by Western blot analysis (Fig.6B). Following two intratumoral administrations of H-1PV, control tumours exhibited a marked reduction in tumour volume (321.5mm^3^±15.9 vs 160mm^3^±27.9, p<0.01) and weight (0.3g±0.04 vs 0.18g±0.02, p<0.05), consistent with robust oncolytic activity, compared to control mice receiving PBS (Fig. 6C and 6D). In contrast, *TGOLN2*-deficient tumours failed to respond to H-1PV treatment, showing no significant reduction in tumour volume or mass (Fig. 6E and 6F). Consistent with this loss of therapeutic efficacy, quantification of viral genomes at the experimental endpoint revealed a significant decrease in H-1PV copy number in *TGOLN2*-depleted tumors compared with controls (4.1×10^3^a.u.±1.7×10^3^ vs 0.3×10^3^a.u.±0.1×10^3^, p<0.05, Fig. 6G). Collectively, these results demonstrate that TGN46 expression is required for productive H-1PV infection and oncolytic activity in experimental PDAC tumour models.

**Figure 6:**
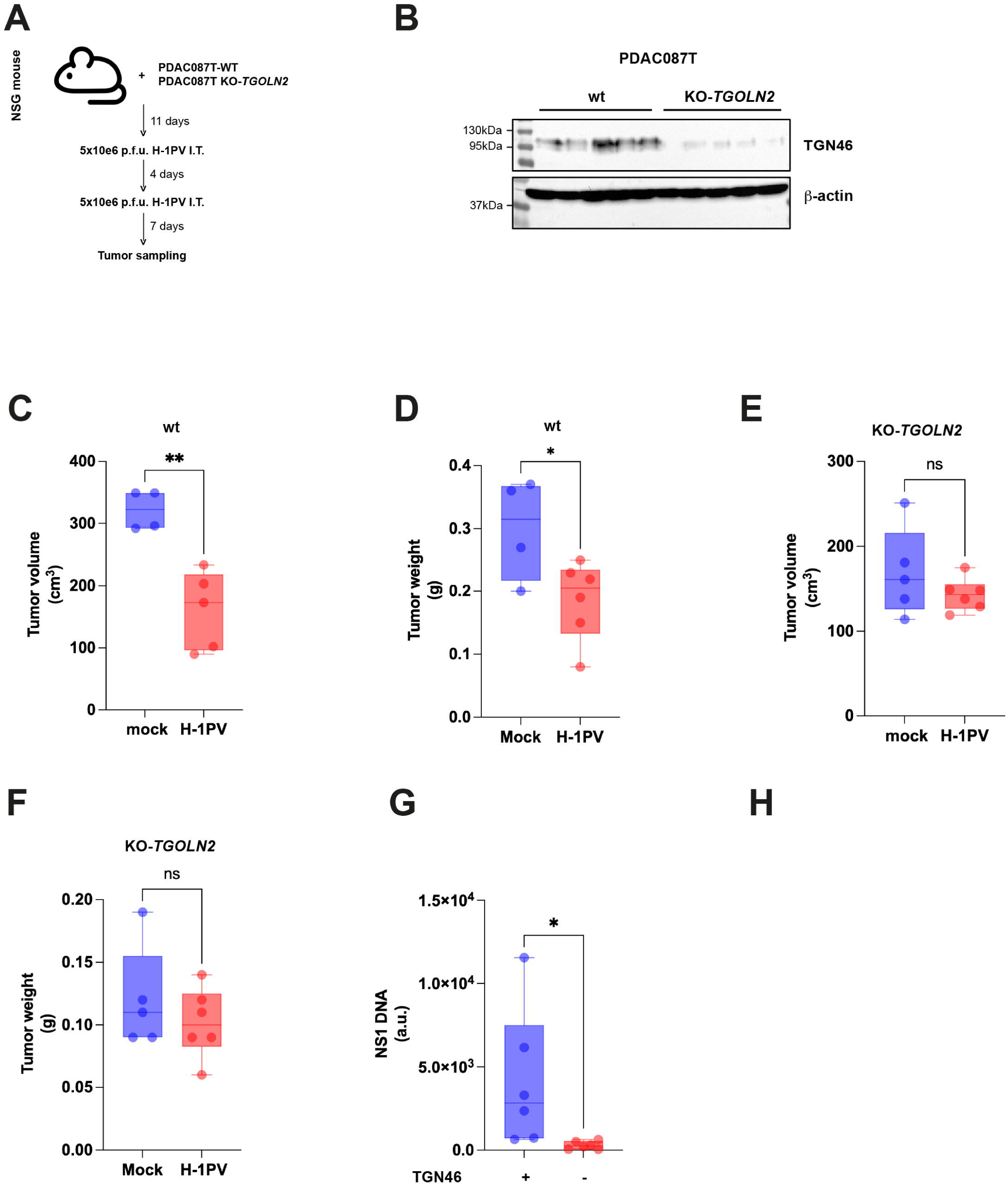
H-1PV antitumoral effects in vivo depend on TGN46. **A**. Schematic representation of the in vivo study. PDAC087T-WT or PDAC087T-*TGOLN2*-KO cells were implanted subcutaneously in NSG mice. Eleven days later, experimental tumors received a first intratumoral (IT) injection of 5 × 10⁶ plaque-forming units (p.f.u.) of the virus, followed by a second injection 4 days later. Control mice received two intratumoral injections of PBS. Tumors were sampled and analyzed 7 days after the second viral injection, and TGN46 expression was assessed by Western blot (**B**). Tumor volume (**C**, **E**), weight (panels **D**, **F**), were measured, and viral content was quantified by NS1 qPCR (**G**). Data represent the mean ± SD of at least 4-6 animals per group. Statistical comparisons were performed using an unpaired t-test.

## Discussion

Oncolytic viruses offer a promising approach to selectively target malignant cells while sparing normal tissues^4^. Among these, H-1PV stands out due to its non-pathogenicity in humans, its natural oncotropism and safety in clinical trials^8,9,10^. However, the molecular basis of this selectivity has remained partly defined. Contrary to many OV, the oncoselectivity of parvoviruses in human cells does not appear to rely on impaired innate immunity within cancer cells. Studies have demonstrated that IFN signalling fails to reduce H-1PV infectivity or progeny production in both normal and transformed cells^34,35^, despite the upregulation of interferon-stimulated genes (ISGs)^35^. This unexpected resistance to IFN-mediated antiviral responses may stem from either the absence of functional pattern recognition receptor (PRR) expression or intrinsic defects, potentially virus-driven, in the endosomal sensing of parvoviral genomes^36^. Notably, H-1PV susceptibility does not correlate with absolute ISG levels but rather with the efficiency of viral entry. This is exemplified by the cooperative role of laminin γ1^33^ and galectin-1^15^, the later recently confirmed as a key mediator of H-1PV efficacy during a biomarker screen in PDAC cell cultures^32^.Yet, despite that H-1PV efficacy in PDAC appears to rely almost entirely on effective cell entry, the identity of high-affinity cell surface receptors facilitating H-1PV uptake has remained elusive, and definitive biomarkers predicting viral susceptibility remains to be fully established.

In this study, we demonstrate that TGN46 is involved early stages of viral entry, supports productive infection, and is required for downstream oncolytic activity. TGN46 is encoded by *TGOLN2* and is widely expressed across various human tissues, particularly in epithelial and secretory cells, where it plays a role in protein sorting and trafficking within the TGN. It functions as a cargo receptor, facilitating the sorting of secretory proteins into specific vesicles for delivery to their destinations^37^. There is no direct link between TGN46 and cancer, but it has been implicated in the formation of CARTS (carriers of the TGN to the cell surface), a PKD-dependent pathway critical for the secretion of specific cargo^38^, particularly that of pancreatic adenocarcinoma upregulated factor (PAUF)^39^, that is associated with TGFβ-dependent increase in the migration and invasion of PDAC cell lines^40^, and possibly in the establishment of an immunosuppressive tumor microenvironment^41^.

Using for the first time in the filed a combination of genetic perturbation, high-resolution imaging, co-immunoprecipitation, and functional virological assays, we demonstrate that TGN46 transiently localizes to the plasma membrane where it engages the H-1PV capsids in Rab5a-enriched domains, prior to CME. This previously underestimated surface function of TGN46 contrasts with the aforementioned classical role as a resident of the TGN, revealing new insights into the dynamic trafficking of this protein and its potential repurposing in the context of viral entry. This echo recent findings indicating that TGN46 plays a role in the intracellular trafficking of SARS-CoV-2 proteins, especially SARS-CoV-2 E protein that undergoes retrograde trafficking to the TGN, accumulating in LAMP1-positive vesicles, likely lysosomes^42^.

Our findings position TGN46 as a *bona fide* host factor for H-1PV infection in PDAC cells and TGN46 and H-1PV virions colocalize at the cell membrane. Mechanistically, TGN46 domain I emerges as both necessary and sufficient for H-1PV internalization following interaction with H-1PV capsid proteins, whereas domains II and III were dispensable. Treatment with sialylation inhibitor neuraminidase or mutation of the single predicted glycosylation site within domain I (N39) impaired binding to VP1 but not VP2, suggesting distinct roles for glycan-mediated interactions in capsid recognition. Interestingly, TGN46 does not bind with the non-structural NS1 protein or galectin-1 itself, suggesting that TGN46 may act as a primary attachment or internalization factor rather than a broader viral scaffold. Despite the strength of these findings, our study has several limitations. First, while we have established a physical association between TGN46 and the H-1PV capsid, the precise molecular interface remains undefined. Structural studies, such as cryo-EM or crosslinking-mass spectrometry, could help map the interaction site and guide rational design of virus mutants with improved binding properties. Second, although TGN46 appears to be expressed and localized to the plasma membrane in PDAC cells, the underlying regulation of expression or trafficking mechanisms are unknown. Future work should determine whether this pattern of expression and localization is driven by oncogenic signalling pathways or specific post-translational modifications, and whether it is unique to PDAC or observed in other H-1PV-sensitive tumors.

Furthermore, our study uncovers several key insights with significant translational potential. First, we quantified H-1PV infection efficiency across a panel of established PDAC cell lines and primary patient-derived tumor cells. Susceptibility varied markedly, with some cell lines showing robust infection and others remaining refractory. Second, we show that TGN46 expression correlates with susceptibility to H-1PV across both cell lines and patient-derived samples. Third, functional studies demonstrate that inducible overexpression of TGN46 in otherwise resistant cells restored susceptibility to infection. In contrast, CRISPR-mediated deletion of *TGOLN2* strongly attenuated viral infectivity and efficacy, both *in vitro* and *in vivo* in experimental tumor models. These data not only validate TGN46 as a host factor for H-1PV, but also indicate that its expression level is rate-limiting for virotherapy efficacy. Collectively, our findings suggest that TGN46 may serve as predictive biomarker of H-1PV responsiveness in PDAC, a notoriously heterogeneous and treatment-refractory malignancy. The ability to stratify PDAC based on TGN46 expression could offer a biomarker-based approach to personalize oncolytic virus therapies. Pre-treatment screening could help identify patients most likely to benefit from virotherapy. Moreover, our data raise the possibility of pharmacologically enhancing TGN46 expression or surface localization to sensitize otherwise refractory tumors. In conclusion, this study reveals an unexpected role for TGN46 in mediating H-1PV entry and positions this protein as a critical determinant of virotherapy responsiveness in PDAC. These insights pave the way for refined biomarker-driven approaches and underscore the value of probing unconventional host-virus interactions in the development of targeted cancer therapeutics.

---

**Figure S1. A.** Schematic representation of the all-in-one plasmid used for CRISPR/Cas9-mediated knockout (KO) of TGN46 (*TGOLN2* sgRNA) by transient transfection followed by ouabain selection. **B.** Western blot analysis of TGN46 expression in PDAC087T-WT and PDAC087T-*TGOLN2*-KO cells. GAPDH was used as loading control. **C.** Immunofluorescence (IF) analysis of PDAC087T-WT (WT) and PDAC087T-*TGOLN2*-KO (KO) cells stained with anti-TGN46 antibody (green) and DAPI for DNA counterstaining (light grey). Images were acquired using a confocal microscope at 63x objective (top row, 20µm scale bars) and regions of interest were magnified 5 times (bottom row). **D.** Western blot analysis of TGN46 expression in Hela cells, and Hela cells expressing *TGOLN2* sgRNA. GAPDH was used as loading control. **E.** Cell viability assay of Hela cells expressing control sgRNA and wild-type for *TGNOL2* (WT), and *TGOLN2*-KO Hela cells (KO) following infection with H-1PV at the indicated MOI. Data represent mean ± S.D. of three independent experiments performed in triplicates. IF analysis of NS1 **(F)** or capsid **(G)** proteins in PDAC087T-WT (WT) and PDAC087T-*TGOLN2*-KO (KO) cells following infection with H-1PV at an MOI of 2.5. Data represent mean ± S.D. of three independent experiments performed in triplicate. Unpaired t-test were performed for statistical analysis. **H.** Schematic representation of all-in-one “pTRE-TGN46” Tet-on lentiviral vector for doxycycline inducible TGN46 overexpression. **I.** Western blot analysis of TGN46 expression in PDAC087T-pTRE-TGN46 cells -/+ dox (TGN46 -/+). HSP90 was used as loading control. **J.** IF of PDAC087T without dox (TGN46 -) and PDAC087T cells overexpressing TGN46 upon dox treatment (TGN46 +), stained with anti-TGN46 antibody (green) and DAPI (light grey). Images were acquired using a confocal microscope at 63X objective (first row, 20µm scale bars) and regions of interest were magnified 5 times. IF analysis of NS1 **(K)** or capsid **(L)** proteins in PDAC087T cells without dox (TGN46 -) and PDAC087T cells overexpressing TGN46 upon dox treatment (TGN46 +), following infection with H-1PV at an MOI of 2.5. Data represent mean ± S.D. of three independent experiments performed in triplicate. Unpaired t-test were performed for statistical analysis.

**Figure S2. A.** PDAC087T-WT (WT) and PDAC087T-*TGOLN2*-KO (KO) cells, and **(B)** PDAC087T cells without doxycycline (TGN46 −) and PDAC087T cells overexpressing TGN46 upon doxycycline treatment (TGN46 +) were stained with antibodies against vesicular transport-related proteins (CHC, Rab5, EEA1, and LAMP1) or analyzed for TFn-647 uptake. Each row shows representative images of staining for one protein of interest (green) or the TFn-647 signal (pink), along with DAPI counterstaining of DNA (light gray). Images were acquired using a confocal microscope with a 63X objective (scale bars: 20 µm). Quantification of total fluorescence intensity, number of objects, and mean object size (µm²) was performed in 30–50 individual cells per condition. Statistical comparisons were conducted using unpaired t tests. PDAC087T-WT cells in interphase **(C)** or prophase **(D)** were stained with antibodies against TGN46 (green) and viral capsids (red), along with DAPI counterstaining of DNA (light gray). Images were acquired using a confocal microscope with a 63× objective (scale bars: 20 µm). PDAC087T-WT cells were treated with hydroxy-dynasore (Dyn) at the indicated concentrations and stained with antibodies against TGN46 (green) or analyzed for TFn-647 uptake (pink), along with DAPI counterstaining of DNA (light gray). **E.** Images were acquired using a confocal microscope with a 20X objective. Transferrin uptake **(F),** TGN46 fluorescence intensity **(G)** and total TGN46-positive area **(H)** were quantified. Data represent mean ± S.D. of three independent experiments performed in triplicate. Šídák’s multiple comparisons test were performed for statistical comparisons. **I.** Representative image and signal quantification showing colocalization of TGN46 (green) and viral capsid (red).

**Figure S3. A.** Schematic representation of TGN46 fusion proteins (FLAG or GFP in N-or C-terminal, red box) and deleted variants (domain I, II or III, doted lines) generated for rescue and GFP-TRAP experiments (subcloning in previously described pTRE inducible lentivector). Legends: signal peptide (SP), luminal part (Lum.), cytoplasmic part (Cyto.) and transmembrane domain (IV). **B.** PDAC087T-WT cells, or PDAC087T-*TGOLN2-*KO cells were transduced with empty control lentiviral vectos (pTRE) or lentivial vectors encoding for TGN46, FLAG-TGN46 and FLAG-TGN46 domain I, II or III deleted mutants. Protein expression was induced by doxycycline (dox), and cells were stained with antibodies against FLAG and fluorescence was quantified. Data represent mean of three independent experiments performed in triplicate. Statistical comparisons were performed using a two-way ANOVA. **C**. PDAC087T-WT cells, or PDAC087T-*TGOLN2-*KO cells were transduced with empty control lentiviral vectos (pTRE) or lentivial vectors encoding for TGN46, EGFP, EGFP-TGN46 and TGN46-EGFP. Proteins expression was induced by doxycycline (dox), and EGFP fluorescence was quantified. Data represent mean of three independent experiments performed in triplicate. Statistical comparisons were performed using a two-way ANOVA. **D**. PDAC087T-WT cells, or PDAC087T-*TGOLN2-*KO cells were transduced with empty control lentiviral vectos (pTRE) or lentivial vectors encoding for TGN46, EGFP, EGFP-TGN46 and TGN46-EGFP. Proteins expression was induced by doxycycline (dox) and infected with H-1PV at an MOI of 2.5 for 24 h. Cells were fixed and stained with an antibody against H-1PV conformational capsid proteins and DAPI for DNA counterstaining (light gray). Images were acquired using a confocal microscope with a 20X objective. Total capsid fluorescence intensity per cell was quantified and normalized to the control condition (red dotted line). Data represent mean ± SD of three independent experiments performed in triplicate. Statistical analysis was performed using uncorrected Fisher’s LSD test. **E**. Schematic representation of the GFP-Trap strategy. F. PDAC087T-*TGOLN2*-KO cells transduced with lentiviral vectors encoding EGFP, EGFP-TGN46, or TGN46-EGFP were infected with H-1PV at an MOI of 2.5. Two days post-infection, cellular proteins were extracted and incubated with GFP-Trap beads. Input, flowthrough and bead-bound proteins were resolved by western blot for EGFP, VP1, VP2, NS1, GAPDH, and galectin-1 (Gal-1). I, input; FT, flowthrough; B, bead-bound fraction. **F**. PDAC087T cells were pre-treated with neuraminidase, then infected with H-1PV at an MOI of 250. Six hours post-infection, cells were fixed and stained with an antibody against the conformational H-1PV capsid protein. Images were acquired using a confocal microscope with a 63X objective, and fluorescence intensity was quantified. Data represent mean ± SD of three independent experiments performed in triplicate. Statistical comparisons were performed using an unpaired t test.

**Figure S4. A.** Western blot analysis of TGN46, galectin-1 (Gal-1), and laminin-γ1 expression in the indicated PDAC-derived cell lines. GAPDH was used as a loading control. Data are representative of three independent experiments. **B**. Western blot analysis of TGN46 expression in wild-type Panc-1 cells (non-targeting control) and Panc-1 cells expressing *TGOLN2* sgRNA. GAPDH was used as a loading control. Data are representative of three independent experiments. **C**. Immunofluorescence analysis of TGN46 expression in AsPC-1 cells either overexpressing (+) or not (–) TGN46. Data represent the mean ± SD of three independent experiments, each performed in triplicate. Statistical comparisons were performed using an unpaired t-test. **D**. Western blot analysis of TGN46 and galectin-1 (Gal-1) expression in the indicated PDAC-derived primary cells. β-actin was used as a loading control. Data are representative of three independent experiments.

## References

1. Wood, L. D., Canto, M. I., Jaffee, E. M. & Simeone, D. M. Pancreatic Cancer: Pathogenesis, Screening, Diagnosis, and Treatment. Gastroenterology 163, 386–402.e1 (2022).

2. Park, W., Chawla, A. & O’Reilly, E. M. Pancreatic Cancer: A Review. JAMA 326, 851–862 (2021).

3. Halbrook, C. J., Lyssiotis, C. A., Pasca di Magliano, M. & Maitra, A. Pancreatic cancer: Advances and challenges. Cell 186, 1729–1754 (2023).

4. Kollmann, C. F., van Montfoort, N., Cordelier, P., Pol, J. & Olagnier, D. Oncolytic virotherapy: Sparking durable anti-tumor immunity through microenvironment modulation. Semin Immunol 80, 101994 (2025).

5. Zhang, T. et al. Talimogene Laherparepvec (T-VEC): A Review of the Recent Advances in Cancer Therapy. J Clin Med 12, 1098 (2023).

6. Quillien, L., Buscail, L. & Cordelier, P. Pancreatic Cancer Cell and Gene Biotherapies: Past, Present, and Future. Hum Gene Ther 34, 150–161 (2023).

7. Runcie, K. et al. Phase I study of intratumoral injection of talimogene laherparepvec for the treatment of advanced pancreatic cancer. Oncologist oyae200 (2024) doi:10.1093/oncolo/oyae200.

8. Angelova, A. L., Geletneky, K., Nüesch, J. P. F. & Rommelaere, J. Tumor Selectivity of Oncolytic Parvoviruses: From in vitro and Animal Models to Cancer Patients. Frontiers in Bioengineering and Biotechnology 3, (2015).

9. Geletneky, K. et al. Oncolytic H-1 Parvovirus Shows Safety and Signs of Immunogenic Activity in a First Phase I/IIa Glioblastoma Trial. Molecular Therapy 25, 2620–2634 (2017).

10. Hajda, J. et al. A non-controlled, single arm, open label, phase II study of intravenous and intratumoral administration of ParvOryx in patients with metastatic, inoperable pancreatic cancer: ParvOryx02 protocol. BMC Cancer 17, 1–11 (2017).

11. Hajda, J. et al. Phase 2 Trial of Oncolytic H-1 Parvovirus Therapy Shows Safety and Signs of Immune System Activation in Patients With Metastatic Pancreatic Ductal Adenocarcinoma. Clinical Cancer Research 27, 5546–5556 (2021).

12. Bretscher, C. & Marchini, A. H-1 Parvovirus as a Cancer-Killing Agent: Past, Present, and Future. Viruses 11, 562 (2019).

13. Hristov, G. et al. Through Its Nonstructural Protein NS1, Parvovirus H-1 Induces Apoptosis via Accumulation of Reactive Oxygen Species. Journal of Virology 84, 5909–5922 (2010).

14. Kulkarni, A. et al. Oncolytic H-1 parvovirus binds to sialic acid on laminins for cell attachment and entry. Nat Commun 12, 3834 (2021).

15. Ferreira, T. et al. Oncolytic H-1 Parvovirus Hijacks Galectin-1 to Enter Cancer Cells. Viruses 14, 1018 (2022).

16. Vasta, G. R. Roles of galectins in infection. Nat Rev Microbiol 7, 424–438 (2009).

17. Ferreira, T. et al. Oncolytic H-1 Parvovirus Enters Cancer Cells through Clathrin-Mediated Endocytosis. Viruses 12, 1199 (2020).

18. Ferreira, T. et al. Oncolytic H-1 Parvovirus Enters Cancer Cells through Clathrin-Mediated Endocytosis. Viruses 12, 1199 (2020).

19. Toolan, H. W., Dalldore, G., Barclay, M., Chandra, S. & Moore, A. E. An unidentified, filtrable agent isolated from transplanted human tumors*. Proceedings of the National Academy of Sciences 46, 1256–1258 (1960).

20. Lujan, P. et al. Sorting of secretory proteins at the trans-Golgi network by TGN46. eLife 12, (2023).

21. Fraunhoffer, N. A. et al. Evidencing a Pancreatic Ductal Adenocarcinoma Subpopulation Sensitive to the Proteasome Inhibitor Carfilzomib. Clin Cancer Res 26, 5506–5519 (2020).

22. Angelova, A. L. et al. Complementary induction of immunogenic cell death by oncolytic parvovirus H-1PV and gemcitabine in pancreatic cancer. J. Virol. 88, 5263–5276 (2014).

23. Joung, J. et al. Genome-scale CRISPR-Cas9 knockout and transcriptional activation screening. Nature Protocols 12, 828–863 (2017).

24. Li, W. et al. MAGeCK enables robust identification of essential genes from genome-scale CRISPR/Cas9 knockout screens. Genome Biology 15, 554 (2014).

25. Agudelo, D. et al. Marker-free coselection for CRISPR-driven genome editing in human cells. Nature Methods 14, 615–620 (2017).

26. Tessmer, C. et al. Generation and Validation of Monoclonal Antibodies Suitable for Detecting and Monitoring Parvovirus Infections. Pathogens 11, 208 (2022).

27. Liu, X. et al. An AP-MS- and BioID-compatible MAC-tag enables comprehensive mapping of protein interactions and subcellular localizations. Nat Commun 9, 1188 (2018).

28. Zhuo, C. et al. Spatiotemporal control of CRISPR/Cas9 gene editing. Signal Transduct Target Ther 6, 238 (2021).

29. Li, J. et al. Synergistic combination of valproic acid and oncolytic parvovirus H-1PV as a potential therapy against cervical and pancreatic carcinomas. EMBO Mol Med 5, 1537–1555 (2013).

30. Mayle, K. M., Le, A. M. & Kamei, D. T. The intracellular trafficking pathway of transferrin. Biochim Biophys Acta 1820, 264–281 (2012).

31. Banting, G., Maile, R. & Roquemore, E. P. The steady state distribution of humTGN46 is not significantly altered in cells defective in clathrin-mediated endocytosis. J Cell Sci 111 **(Pt** **23**), 3451–3458 (1998).

32. Schäfer, T. E. et al. Biomarker screen for efficacy of oncolytic virotherapy in patient-derived pancreatic cancer cultures. EBioMedicine 105, 105219 (2024).

33. Kulkarni, A. et al. Oncolytic H-1 parvovirus binds to sialic acid on laminins for cell attachment and entry. Nat Commun 12, 3834 (2021).

34. Paglino, J. C., Andres, W. & van den Pol, A. N. Autonomous parvoviruses neither stimulate nor are inhibited by the type I interferon response in human normal or cancer cells. J Virol 88, 4932–4942 (2014).

35. Neulinger-Muñoz, M. et al. Human Retrotransposons and the Global Shutdown of Homeostatic Innate Immunity by Oncolytic Parvovirus H-1PV in Pancreatic Cancer. Viruses 13, 1019 (2021).

36. Raykov, Z. et al. TLR-9 contributes to the antiviral innate immune sensing of rodent parvoviruses MVMp and H-1PV by normal human immune cells. PLoS One 8, e55086 (2013).

37. Lujan, P. et al. Sorting of secretory proteins at the trans-Golgi network by human TGN46. eLife 12, RP91708 (2024).

38. Wakana, Y. et al. Kinesin-5/Eg5 is important for transport of CARTS from the trans-Golgi network to the cell surface. Journal of Cell Biology 202, 241–250 (2013).

39. Wakana, Y. et al. A new class of carriers that transport selective cargo from the trans Golgi network to the cell surface. The EMBO Journal 31, 3976–3990 (2012).

40. Lee, M. et al. TGF-β-Induced PAUF Plays a Pivotal Role in the Migration and Invasion of Human Pancreatic Ductal Adenocarcinoma Cell Line Panc-1. Int J Mol Sci 25, 11420 (2024).

41. Kim, Y. J. et al. Pancreatic Adenocarcinoma Up-Regulated Factor (PAUF) Transforms Human Monocytes into Alternative M2 Macrophages with Immunosuppressive Action. Int J Mol Sci 25, 11545 (2024).

42. Hutchison, J. M. et al. Recombinant SARS-CoV-2 envelope protein traffics to the trans-Golgi network following amphipol-mediated delivery into human cells. J Biol Chem 297, 100940 (2021).

43. Hatoyama, Y., Homma, Y., Hiragi, S. & Fukuda, M. Establishment and analysis of conditional Rab1- and Rab5-knockout cells using the auxin-inducible degron system. J Cell Sci 134, jcs259184 (2021).

44. McLauchlan, H. et al. A novel role for Rab5-GDI in ligand sequestration into clathrin-coated pits. Curr Biol 8, 34–45 (1998).

