## Supplemental figures for "A genome-wide CRISPR-Cas9 screen identifies TGN46 as a host determinant for H-1PV susceptibility in pancreatic adenocarcinoma"

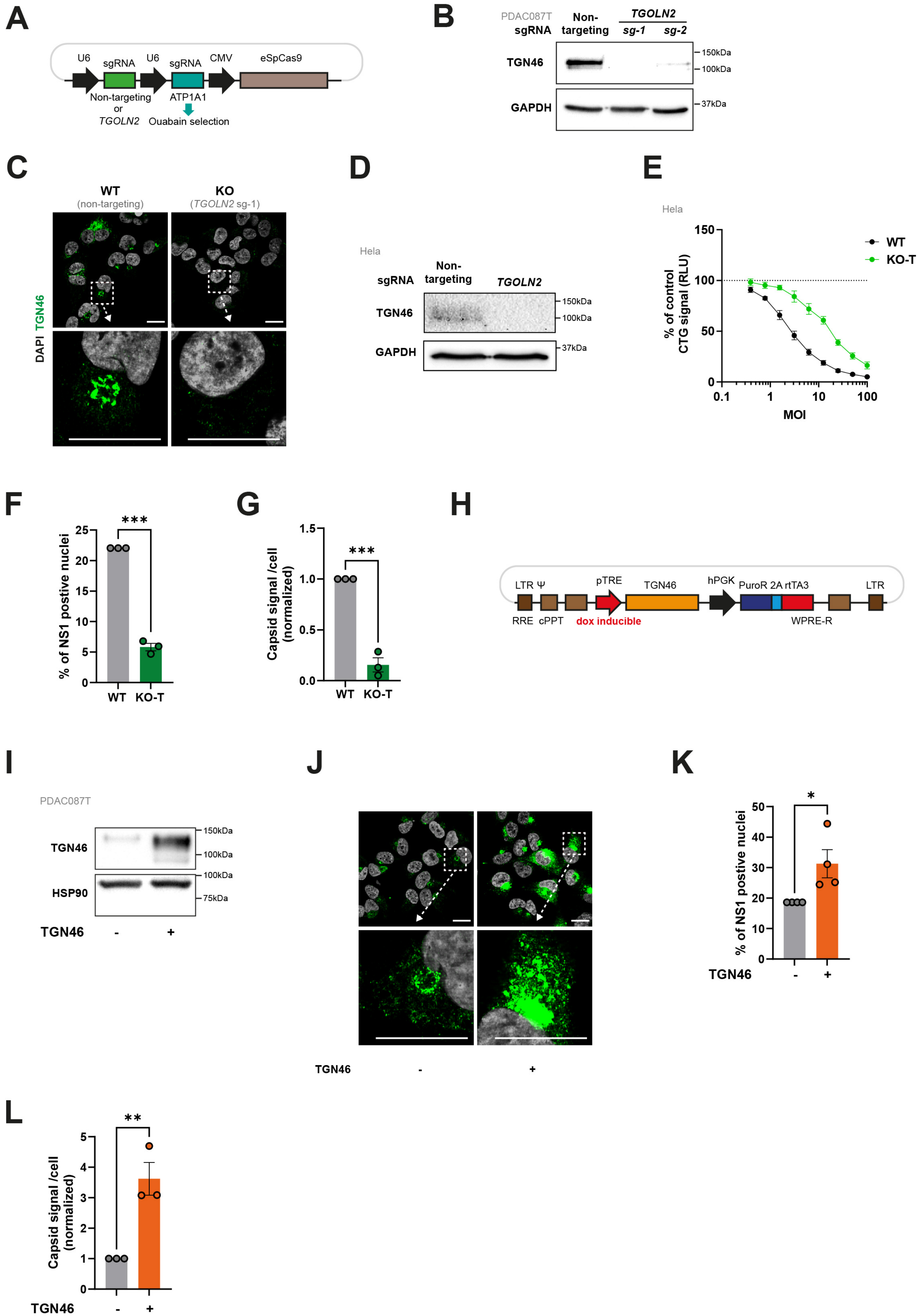

A
U6
sgRNA
U6
sgRNA
CMV
eSpCas9
Non-targeting
or
TGOLN2
ATP1A1
Ouabain selection
B
PDAC087T
sgRNA
Non-targeting
TGOLN2
sg-1
sg-2
TGN46
GAPDH
150kDa
100kDa
37kDa
C
WT
(non-targeting)
KO
(TGOLN2 sg-1)
DAPI TGN46
D
Hela
sgRNA
Non-targeting
TGOLN2
TGN46
GAPDH
150kDa
100kDa
37kDa
E
Hela
% of control
CTG signal (RLU)
WT
KO-T
MOI
F
% of NS1 postive nuclei
***
WT
KO-T
G
Capsid signal /cell
(normalized)
***
WT
KO-T
H
LTR Ψ
RRE
cPPT
pTRE
dox inducible
TGN46
hPGK
PuroR 2A rtTA3
WPRE-R
LTR
I
PDAC087T
TGN46
HSP90
150kDa
100kDa
100kDa
75kDa
TGN46
J
TGN46
K
% of NS1 postive nuclei
*
TGN46
L
Capsid signal /cell
(normalized)
**
TGN46

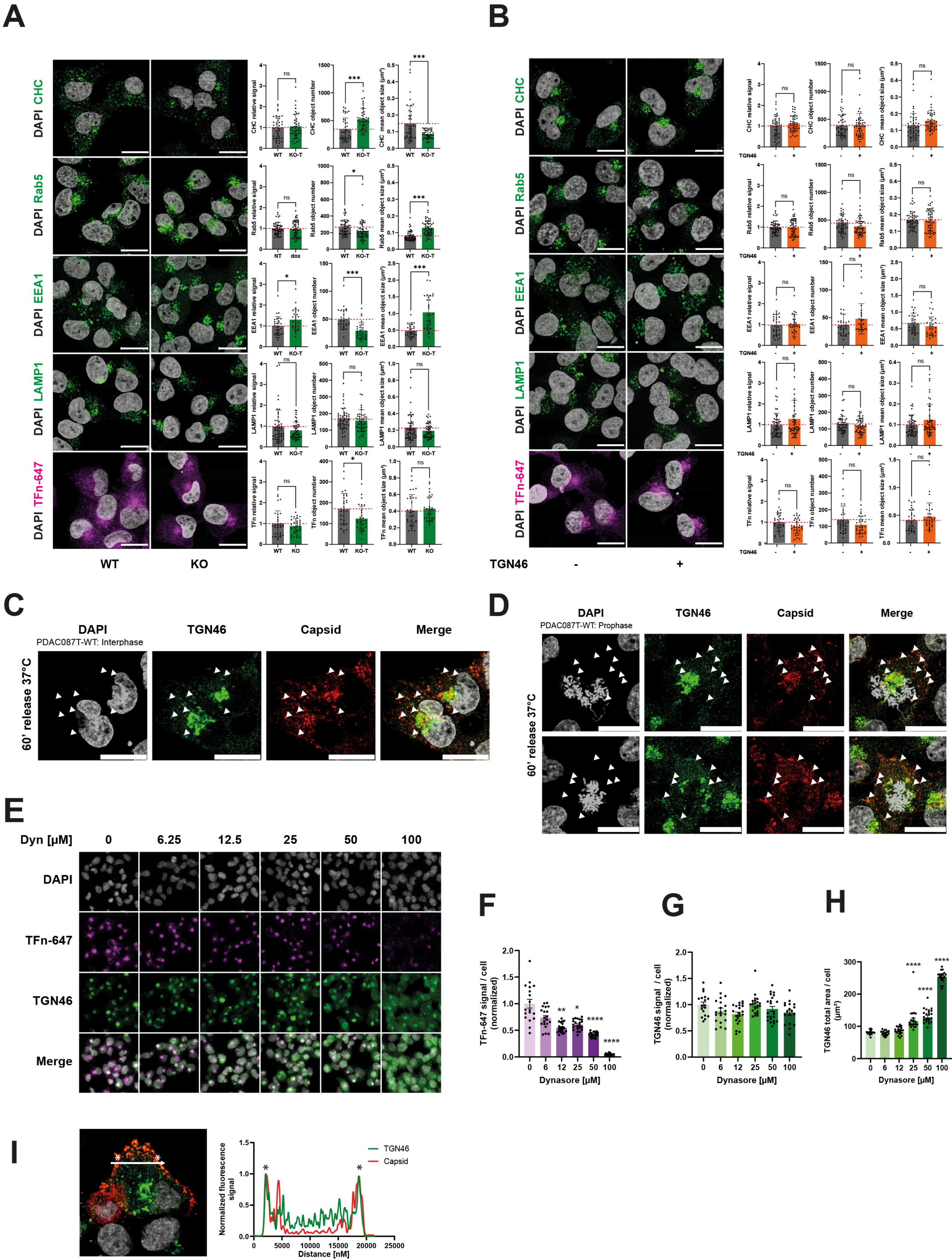
A
B
DAPI CHC
DAPI Rab5
DAPI EEA1
DAPI LAMP1
DAPI TFn-647
WT
KO
CHC relative signal
CHC object number
CHC mean object size (µm²)
Rab5 relative signal
Rab5 object number
Rab5 mean object size (µm²)
EEA1 relative signal
EEA1 object number
EEA1 mean object size (µm²)
LAMP1 relative signal
LAMP1 object number
LAMP1 mean object size (µm²)
TFn relative signal
TFn object number
TFn mean object size (µm²)
WT KO-T
NT dox
ns
*
***
TGN46
-
+
C
DAPI
TGN46
Capsid
Merge
PDAC087T-WT: Interphase
60' release 37°C
D
PDAC087T-WT: Prophase
E
Dyn [µM]
0
6.25
12.5
25
50
100
TFn-647
Merge
F
G
H
TFn-647 signal / cell (normalized)
TGN46 signal / cell (normalized)
TGN46 total area / cell (µm²)
Dynasore [µM]
**
****
I
Normalized fluorescence signal
Distance [nM]
TGN46
Capsid

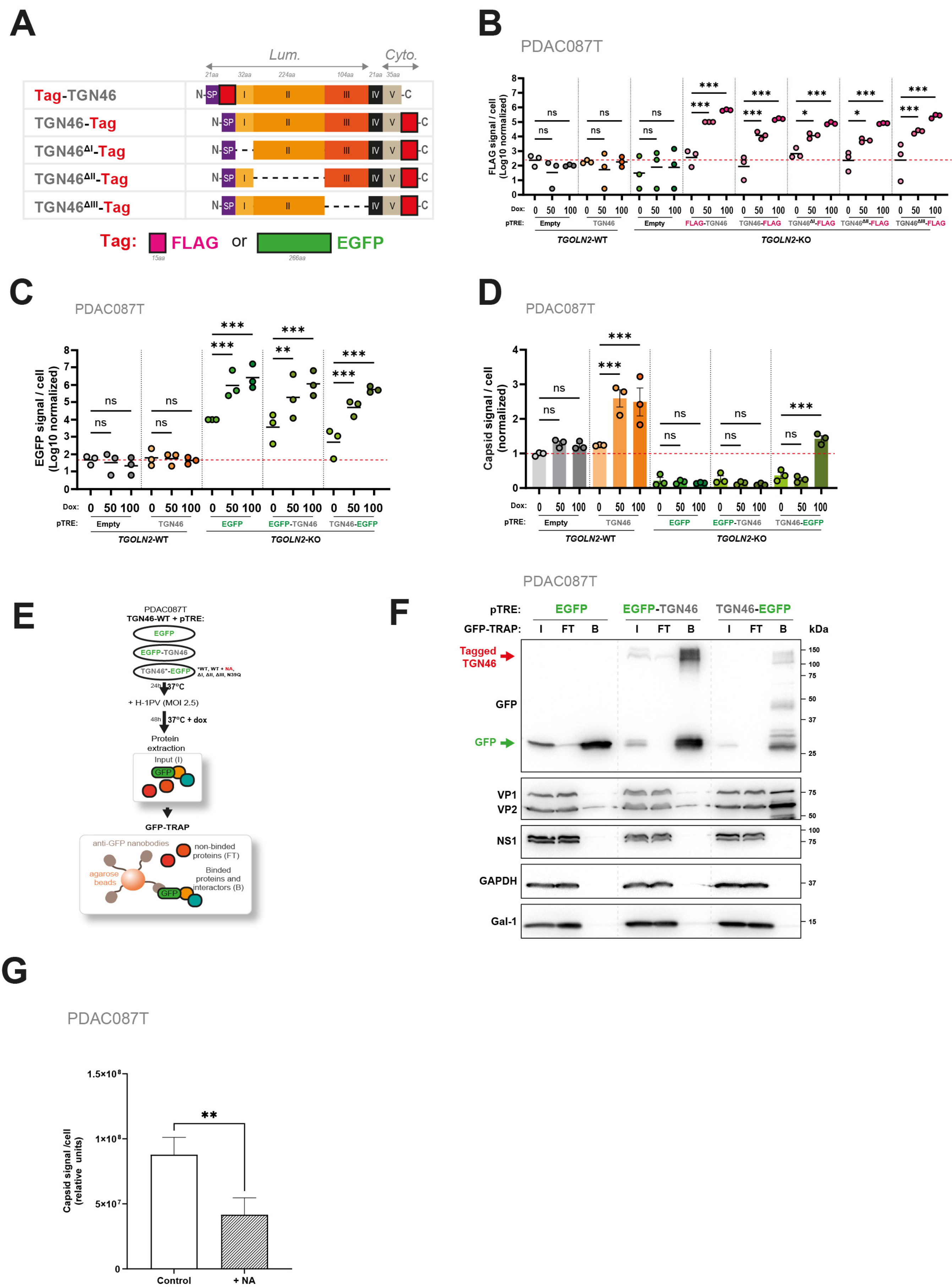
A
Lum.
Cyto.
21aa 32aa 224aa 104aa 21aa 35aa
Tag-TGN46
TGN46-Tag
TGN46ΔI-Tag
TGN46ΔII-Tag
TGN46ΔIII-Tag
N- SP I II III IV V -C
Tag: FLAG or EGFP
15aa 266aa
B
PDAC087T
FLAG signal / cell (Log10 normalized)
ns
***
**
*
Dox: 0 50 100
pTRE: Empty TGN46 Empty FLAG-TGN46 TGN46-FLAG TGN46ΔI-FLAG TGN46ΔII-FLAG TGN46ΔIII-FLAG
TGOLN2-WT
TGOLN2-KO
C
PDAC087T
EGFP signal / cell (Log10 normalized)
Dox: 0 50 100
pTRE: Empty TGN46 EGFP EGFP-TGN46 TGN46-EGFP
TGOLN2-WT
TGOLN2-KO
D
PDAC087T
Capsid signal / cell (normalized)
Dox: 0 50 100
pTRE: Empty TGN46 EGFP EGFP-TGN46 TGN46-EGFP
TGOLN2-WT
TGOLN2-KO
E
PDAC087T
TGN46-WT + pTRE:
EGFP
EGFP-TGN46
TGN46*-EGFP
*WT, WT + NA, ΔI, ΔII, ΔIII, N39Q
24h 37°C
+ H-1PV (MOI 2.5)
48h 37°C + dox
Protein extraction
Input (I)
GFP
GFP-TRAP
anti-GFP nanobodies
non-binded proteins (FT)
agarose beads
Binded proteins and interactors (B)
F
PDAC087T
pTRE: EGFP EGFP-TGN46 TGN46-EGFP
GFP-TRAP: I FT B I FT B I FT B
kDa
Tagged TGN46
GFP
GFP
150 100 75 50 37 25
VP1
VP2
NS1
GAPDH
Gal-1
15
G
PDAC087T
Capsid signal /cell (relative units)
1.5x10⁸
1x10⁸
5x10⁷
0
**
Control
+ NA

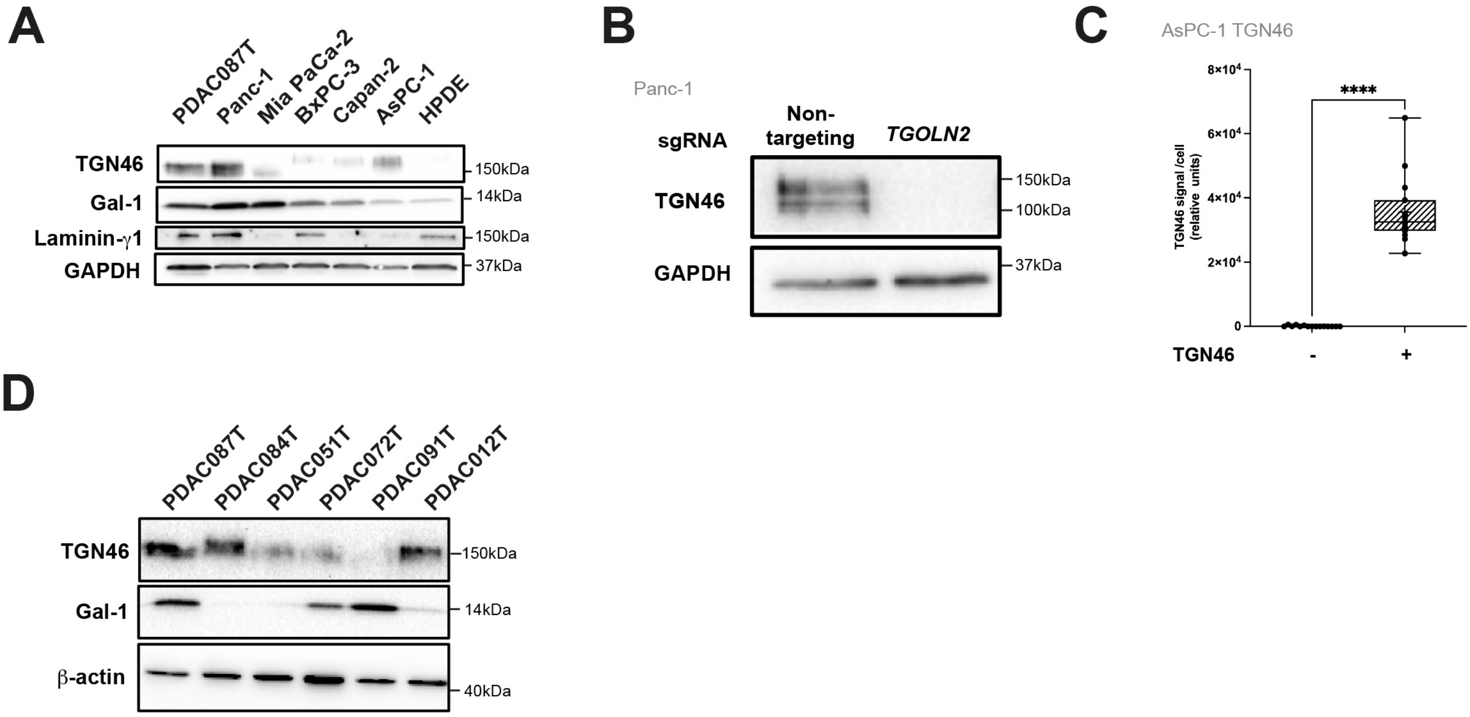
A
PDAC087T
Panc-1
Mia PaCa-2
BxPC-3
Capan-2
AsPC-1
HPDE
TGN46
Gal-1
Laminin-γ1
GAPDH
150kDa
14kDa
150kDa
37kDa
B
Panc-1
sgRNA
Non-targeting
TGOLN2
TGN46
GAPDH
150kDa
100kDa
37kDa
C
AsPC-1 TGN46
TGN46 signal /cell (relative units)
8×10^4
6×10^4
4×10^4
2×10^4
0
****
TGN46
-
+
D
PDAC087T
PDAC084T
PDAC051T
PDAC072T
PDAC091T
PDAC012T
TGN46
Gal-1
β-actin
150kDa
14kDa
40kDa

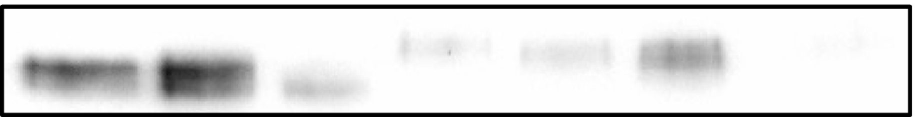

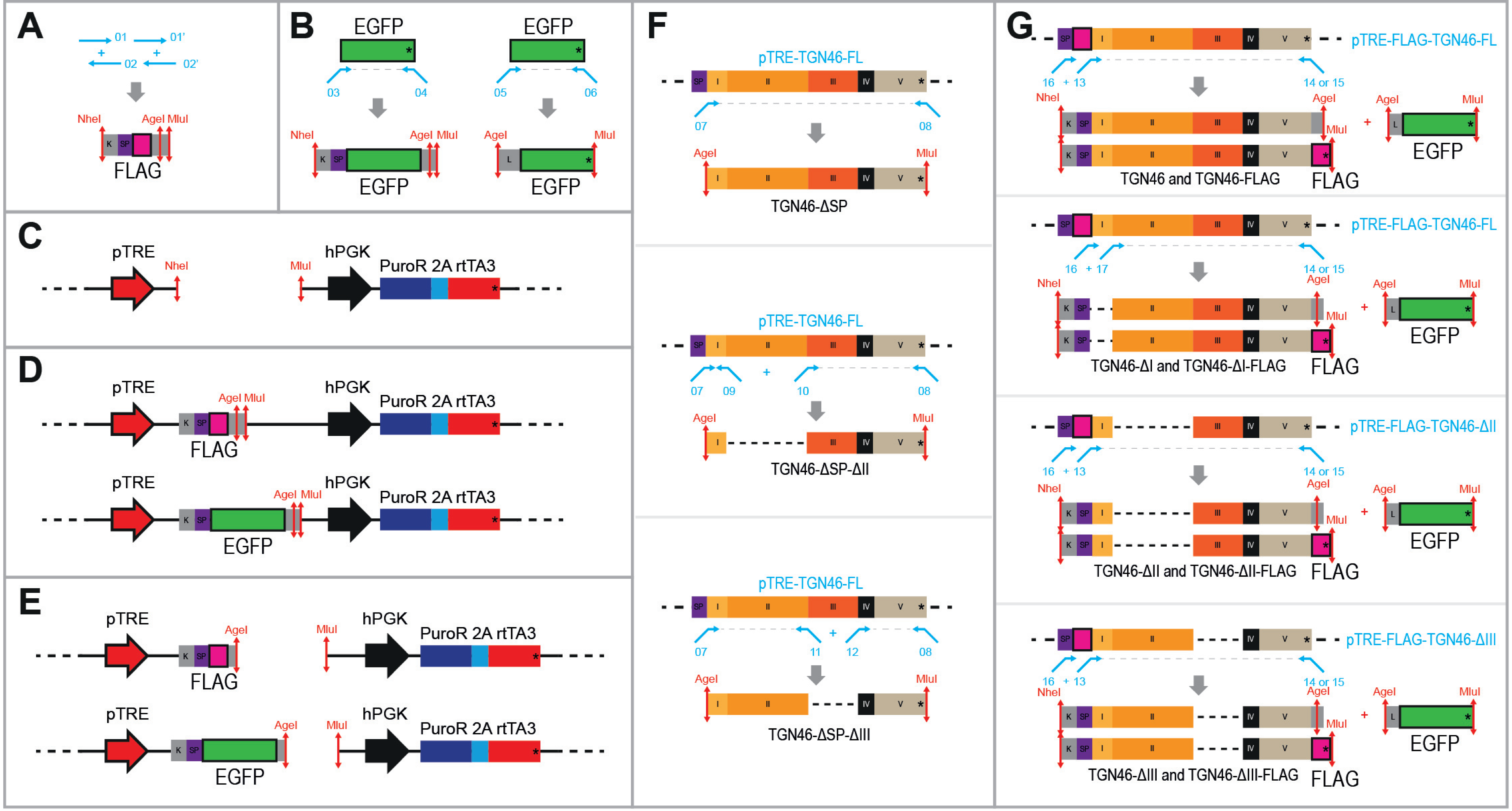


**Illustration 1. Schematic representation of *TGOLN2* mutagenesis and fusion with EGFP and FLAG tags.** Amplification of FLAG (**A**) and EGFP (**B**) for cloning in pTRE-hPGK-PuroR2ArtTA3 expression vector. **C**. Digestion of pTRE-hPGK-PuroR2ArtTA3 with Nhe1 and MluI restriction enzymes. **D-E**. Cloning of FLAG or EGFP into pTRE-hPGK-PuroR2ArtTA3 expression vector. **F**. PCR amplification of *TGOLN2* deleted for the signal peptide (ΔSP) and domain II (ΔII) or III (ΔIII) from pTRE-*TGOLN2*-full lenght (FL). **G**. Cloning by PCR of *TGOLN2* EGFP or *TGOLN*2 FLAG, wild-type or deleted from domain 1 (ΔI), 2 (ΔII) or 3 (ΔIII) using pTRE-*TGOLN2*-full lenght (FL) as a matrix.
